# Cellular genome wide association study identifies common genetic variation influencing lithium induced neural progenitor proliferation

**DOI:** 10.1101/2022.01.31.478307

**Authors:** Justin M. Wolter, Brandon D. Le, Nana Matoba, Michael J. Lafferty, Nil Aygün, Dan Liang, Kenan Courtney, Joseph Piven, Mark J. Zylka, Jason L. Stein

## Abstract

Lithium is used in the treatment of bipolar disorder (BD) and is known to increase neural progenitor cell (NPC) proliferation. Though the mechanism of lithium’s therapeutic effect is not understood, evidence suggests that genetic variation influences response to treatment. Here, we used a library of genetically diverse human NPCs to identify common genetic variants that modulate lithium induced proliferation. We identified a locus on chr3p21.1 associated with lithium induced proliferation that colocalizes with BD risk. One lithium responsive gene, *GNL3*, was detected within the locus. The allele associated with increased baseline and lithium-induced *GNL3* expression was also associated with increased lithium-induced NPC proliferation. Experimental manipulation of *GNL3* expression using CRISPRa/i in NPCs showed that *GNL3* was necessary for lithium’s full proliferative effects, and sufficient to induce proliferation without lithium treatment. In all, our data suggest that *GNL3* expression sensitizes NPCs for a stronger proliferative response to lithium.

## Main Text

Bipolar disorder (BD) is highly heritable and commonly treated using lithium salts^1–4^. However, only ∼40-60% of individuals with BD who are treated with lithium respond favorably^5–7^. Genetic background influences lithium’s efficacy^8^, as demonstrated by genome wide association studies (GWAS) of lithium response in individuals with BD^9,10^, and that iPSC lines generated from individuals with BD recapitulate lithium responsiveness^11–14^. Despite recent advances in identifying the genetic loci that underlie BD risk^4^, much less is known of the genetic underpinnings of lithium responsiveness^15^. This is at least in part due to the difficulty of performing large pharmacogenomic studies in human populations, and the context-dependent effects of lithium on diverse cell types^16^. Though the mechanism underlying lithium’s therapeutic effects in BD are unknown, lithium induces proliferation in adult hippocampal NPCs^24,25^, an effect required for the therapeutic action of some antidepressant drugs^26–29^. Lithium is classified as a class D teratogen, and usage is generally discouraged during pregnancy except in cases where concerns over relapse of bipolar episodes outweigh increased risk of fetal cardiac malformations^17,18^. Teratogenic effects on human brain tissues resulting from fetal exposure to lithium are controversial^19^, and while subtle neurodevelopmental alterations have been reported in animal models^20–23^, whether fetal NPC proliferation contributes to these effects is unknown.

Based on these lines of evidence we sought to identify genetic variants influencing lithium-induced proliferation in primary NPCs derived from a population of genetically diverse donors which have previously been extensively characterized by genotyping, RNA-seq, and ATAC-seq^30–32^. We measured the effect of LiCl on NPC proliferation using an EdU incorporation assay, which fluorescently labeled cells in S-phase of the cell cycle at any point during a two hour EdU pulse. Following the EdU pulse, cells were fixed, additionally labeled with a fluorescent DNA content dye, and the number of cells within each phase of the cell cycle was quantified using flow cytometry (Fig. 1a). We found that lithium increased NPC proliferation, as quantified by the percentage of cells in S-phase, at relatively low concentrations (≤2.5 mM), and decreased proliferation at higher concentrations (≥5 mM) (Extended Data Fig. 1a,b). Lithium has been shown to induce NPC proliferation by activating Wnt signaling through competitive inhibition of GSK3B^33^, but we did not observe activated Wnt signaling at proliferation-inducing concentrations using either a beta catenin activated luciferase reporter assay or endogenous Wnt target gene expression (Extended Data Fig. 1c-e). GSK3B directly regulates many transcription factors which in turn activate signaling pathways such as CREB and NFAT^34^. We found that lithium activated expression of target genes of the NFAT signaling pathway at proliferation-inducing concentrations in NPCs^35^, but did not detectably affect gene expression of Wnt or CREB pathway associated genes (Extended Data Fig. 1g-i), suggesting that lithium induced NPC proliferation was not mediated by Wnt pathway activation.

**Figure 1:**
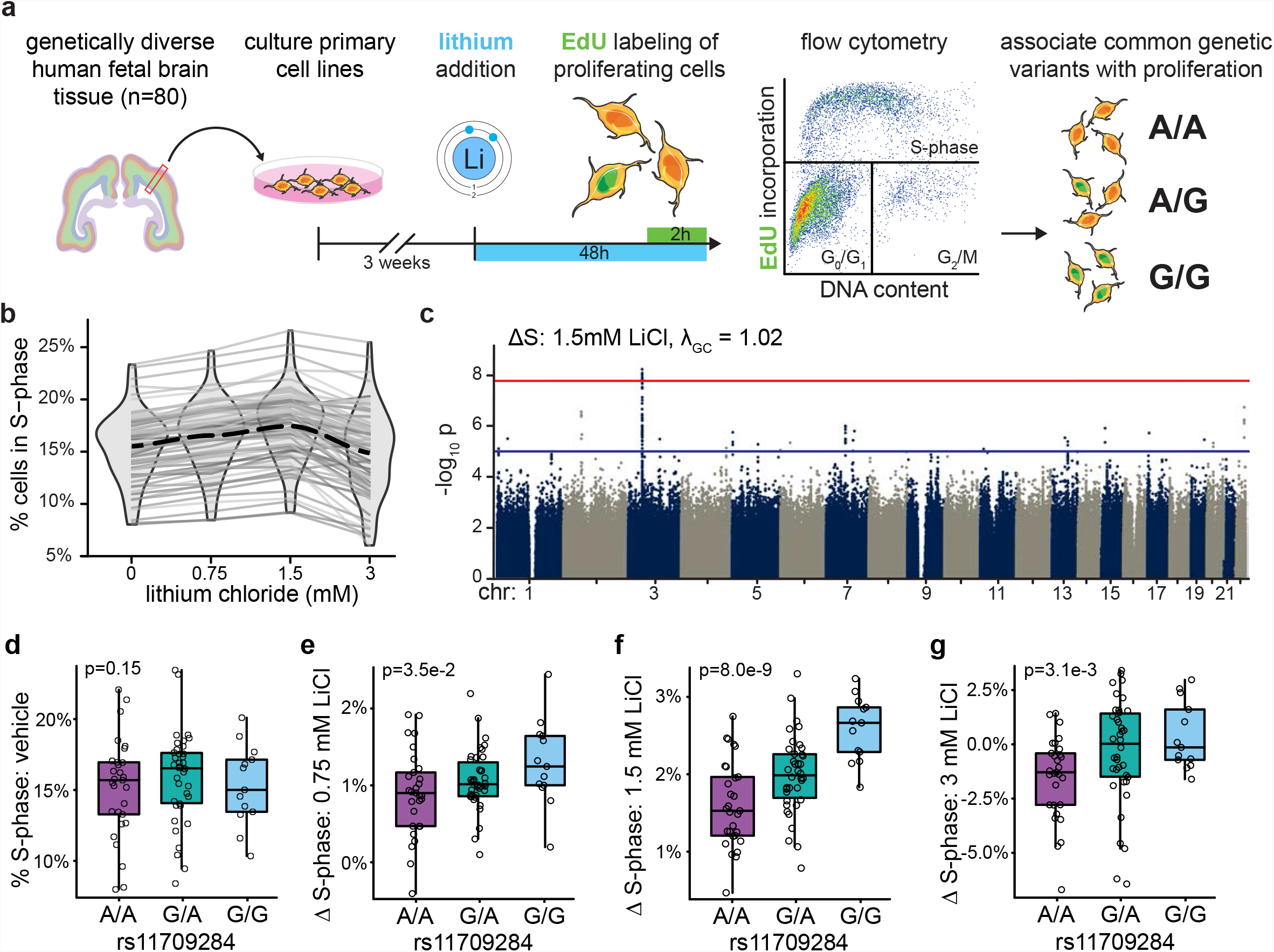
Cellular GWAS identifies genomic loci associated with lithium induced proliferation in NPCs. **a)** Experimental strategy to identify common genetic variants associated with lithium-induced NPC proliferation. **b)** Percentage of cells in S-phase measured by EdU incorporation across indicated concentrations of LiCl in n=80 distinct NPC donors, each represented by a gray line. The mean across donors is represented by the black dotted line. The distribution of cells in S-phase for each concentration is shown as a violin plot. **c)** Manhattan plot showing GWAS of ΔS-phase in response to 1.5 mM LiCl. The red line denotes study-wide significance threshold (p = 1.67 × 10^−8^). The blue line denotes nominal significance threshold (p = 1 × 10^−5^). **d-g)** Boxplots showing proliferation across genotypes at SNP rs11709284 in response to vehicle (**d**), ΔS-phase at 0.75mM (**e**), 1.5mM (**f**), and 3mM LiCl (**g**). Each point is a single NPC donor line with the indicated genotype. For all boxplots, the center line represents the median value, the bounds of the box are 25th and 75th percentiles, and the whiskers equal 1.5 times the interquartile range.

As the target serum concentration for long term lithium treatment is 0.6-1.2 mM, we focused on concentrations that approximate therapeutically relevant levels (0.75 and 1.5mM), as well as a higher concentration (3mM) that we hypothesized may be associated with negative side effects^36^. We measured lithium-induced proliferation using the previously described EdU incorporation and flow cytometry assay in a population of n=80 genetically diverse NPC donors (Fig. 1a, see Extended Data Fig. 2, 3a-c and Methods). NPCs from all donor lines increased proliferation in a concentration dependent manner at 0.75 and 1.5 mM LiCl, while some NPC donor lines decreased proliferation at 3mM (Fig. 1b). Phenotypic values were normally distributed, and technical replicates (the same donor thawed at different times) were reasonably correlated (Extended data Fig. 3d-h). Proliferation rates in vehicle and lithium conditions were strongly correlated (Extended Data Fig. 3i), which could mask the effect of genetic variation on lithium-induced proliferation. Therefore, our primary phenotype, termed ΔS-phase, subtracted the vehicle proliferation rate from the lithium exposure proliferation rate.

To identify genetic variants associated with lithium-induced NPC proliferation, we performed genome-wide association tests using a linear mixed effects model^37^. While proliferation rates were not correlated with any measured biological or technical variables (Extended Data Figs. 3j-n), we included gestation week, sex, and multi-dimensional scaling components to control for ancestry as covariates in the association model. We identified 80 nominally significant (p < 1×10^−5^) loci across all conditions (vehicle, ΔS-phase 0.75, 1.5, and 3 mM LiCl), and one study-wide significant association (p < 1.67×10^−8^, corrected for the number of independent tests^38^) (Fig. 1c, Extended Data Fig. 4, Supplemental Table 1). The study-wide significant locus was associated with lithium induced proliferation (ΔS-phase) at 1.5 mM LiCl, but the association was attenuated in response to other lithium concentrations, and not detectably associated with vehicle proliferation rate (Fig. 1d-g). In all, these results demonstrate that GWAS performed in cultured cells can identify novel context-dependent pharmacogenomic interactions^39^.

The study-wide significant locus was found on chr3p21.1, and spans a >500kb gene dense region (Fig. 2a). This locus contains two SNPs with the lowest study-wide p-values (rs352140, p = 5.71×10^−9^; rs11709284, p = 8.00×10^−9^), which are weakly correlated by linkage disequilibrium (LD, *r*^*2*^ = 0.22). To evaluate whether rs352140 and rs11709284 tag independent loci, we conditioned the associations on either SNP, both of which reduced the significance of the association of the other, indicating that the association signal identifies a single but complex locus (Extended Data Fig. 5a-c).

**Figure 2:**
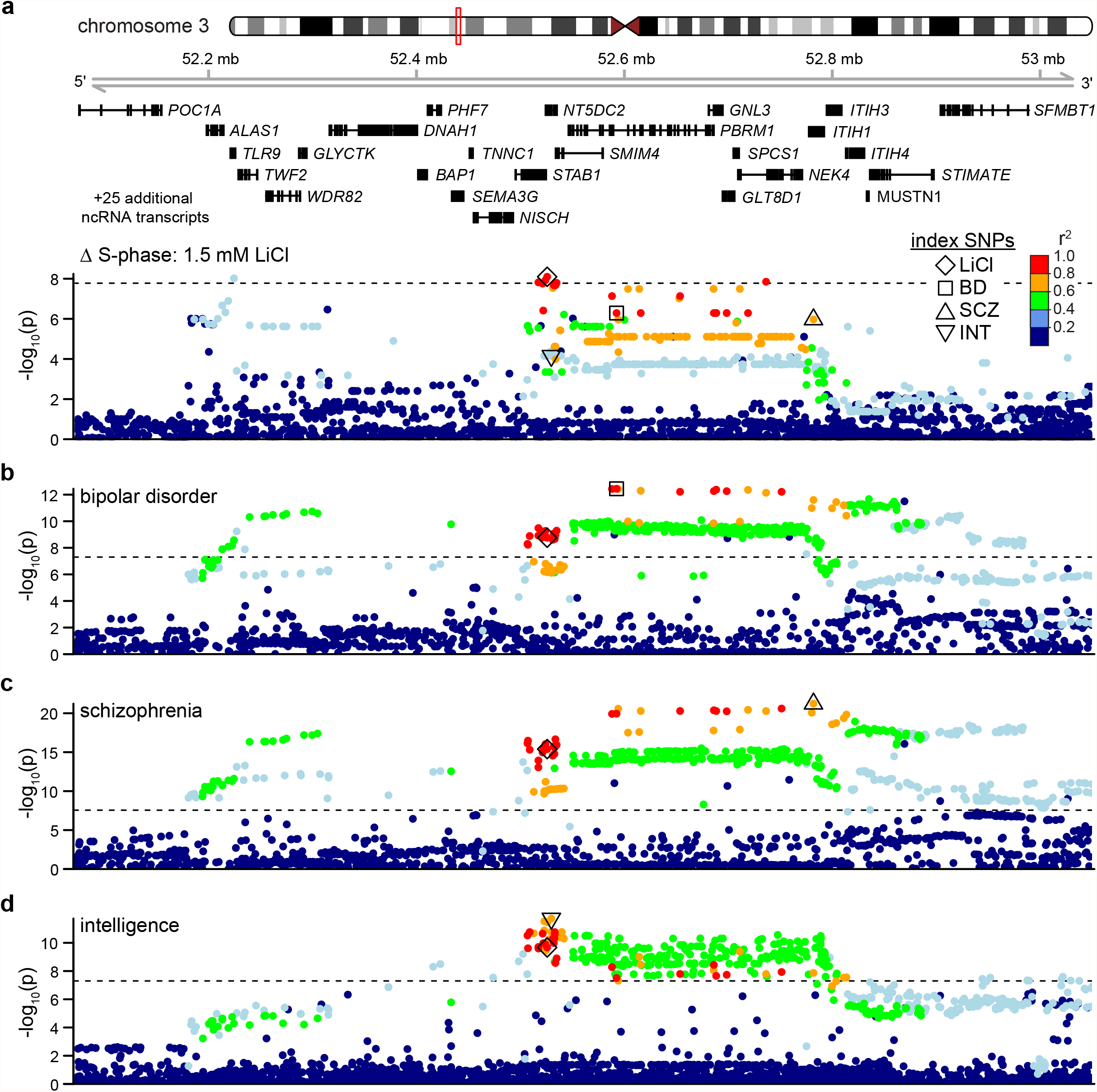
Lithium-induced proliferation at chr3p21.1 colocalizes with neuropsychiatric disorders and intelligence GWAS. **a)** Locus zoom of the 1.5 mM LiCl ΔS-phase phenotype. Dashed line denotes study-wide significance level. ◊: SNP of interest from this study (rs11709284). Each SNP is colored by LD (*r*^*2*^) to rs11709284 in NPC donors **b-d**) GWAS results for BD^4^ (**b**), schizophrenia^41^ (**c**), and intelligence^42^ (**d**) at the same locus as (a). SNP annotations: ◻: index SNP for BD (rs2336147). △: index SNP for schizophrenia (rs2710323). ▽: index SNP for intelligence (rs4687625). *r*^*2*^ values relative to rs11709284 were calculated using the 1000 Genomes EUR reference panel^61^. Dashed lines indicate standard genome wide significance threshold (p = 5 × 10^−8^).

To explore the association of this locus to other complex brain traits we tested for colocalization with existing brain-related GWAS summary statistics using conditional analysis^40^. We identified colocalizations with risk for BD^4^, schizophrenia^41^, and inter-individual differences in intelligence^42^ (Fig. 2b-d). Conditioning on each index SNP from these brain-related GWAS markedly reduced proliferation associations, especially for rs11709284 (Extended Data Fig. 5d-f), suggesting that causal variant(s) affecting lithium-induced proliferation and these complex brain traits are shared. We focus on rs11709284 as marking the lithium-induced proliferation associated locus given its stronger association with these other traits. The lithium-induced proliferation-increasing allele is a risk allele for BD and schizophrenia, and an intelligence-decreasing allele. The lithium induced proliferation locus was not associated with lithium response in individuals with BD^9^ (Extended Data Fig. 5g), nor was the lone genome-wide significant locus associated with lithium response in individuals with BD associated with lithium induced proliferation or risk for BD (Extended Data Fig. 6). To explore patterns of polygenic overlap between lithium-sensitive proliferation and other traits, we selected variants associated with 1.5mM lithium-induced proliferation at increasing significance thresholds and plotted other trait GWAS p-values at these selected variants using quantile-quantile (QQ) plots (Extended Data Fig 7, Supplemental Table 2). SNPs most significantly associated with lithium-induced NPC proliferation phenotype showed highest enrichment in neuropsychiatric disorder GWAS (bipolar disorder, schizophrenia, major depressive disorder, autism spectrum disorder), relatively moderate enrichment in non-disorder brain traits (intelligence, global cortical thickness), and little enrichment in non-brain traits (HDL being the only exception) (Extended Data Fig. 7a-d). This pattern of enrichment suggests that the genetic variants associated with lithium-sensitive NPC proliferation are also associated with risk for neuropsychiatric disorders.

To identify putative causal genes underlying the association with lithium induced proliferation, we considered a ∼1.2 Mb window containing 29 protein coding genes surrounding rs11709284 (Extended Data Fig. 8a). Previous expression measurements via RNA-seq in these NPCs showed that 21 of these genes had detectable expression at baseline^30,31^ (Extended Data Fig. 8b). We quantified the expression of each of these genes over time in response to 1.5 mM LiCl in one NPC donor line using qPCR, and *GNL3* was the only gene with significantly different expression at multiple timepoints (Extended Data Fig. 8c). Repeating this experiment in NPCs from three additional donors confirmed that 1.5 mM LiCl transiently induces *GNL3* expression, peaking eight hours post exposure (Fig. 3a). Increasing lithium concentration also resulted in relatively higher *GNL3* expression at 8 hours, without extending the duration of expression (Extended Data Fig. 8d).

**Figure 3:**
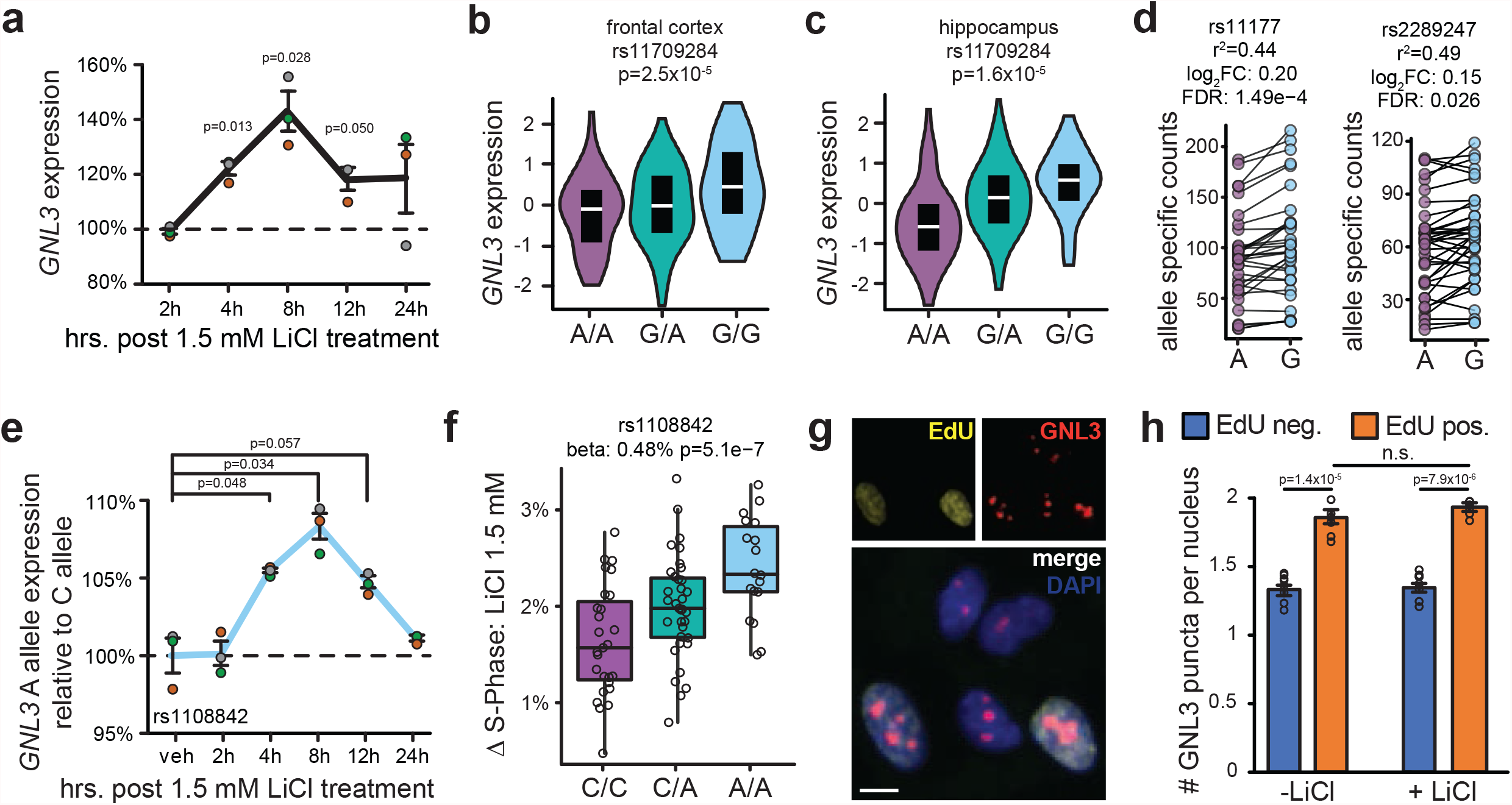
Lithium induced proliferation increasing alleles are also *GNL3* increasing alleles. **a)** *GNL3* expression in NPCs from three randomly selected donor lines following treatment with 1.5 mM LiCl, quantified by RT-qPCR. Each time-point was normalized to vehicle-treated samples extracted at the same time (dashed line). Each colored dot is a distinct NPC donor line. All error bars throughout this study are standard error of the mean. **b**,**c)** Allelic effects on *GNL3* expression at SNP rs11709284 in frontal cortex and hippocampus from GTEx^46^, FDR < 0.05. Alleles associated with increased *GNL3* expression are also associated with increased ΔS-phase (compare alleles with Fig. 1f). In all subsequent figures purple denotes the allele associated with less proliferation, whereas blue denotes the allele associated with increased proliferation from this study. **d)** Allele specific expression of two SNPs in *GNL3* open reading frame in NPCs where donor lines are heterozygous for each SNP, assessed from baseline RNAseq data^30^. *r*^*2*^ values relative to rs11709284. Alleles associated with increased *GNL3* expression at baseline are also associated with increased lithium ΔS-phase (compare alleles with Extended Data Fig. 9h). **e)** Allele specific RT-qPCR for *GNL3* in response to 1.5 mM LiCl, quantified in NPCs from n=3 distinct donors which are heterozygous for a SNP in *GNL3* 5’-UTR (rs1108842). Relative expression of each allele was quantified by calculating the cycle difference between each allele at 0.5 Rn, as in^60^. Each colored dot represents NPCs from a distinct donor. Four technical qPCR replicates were acquired per donor and averaged, and the data was then normalized to vehicle, as in Fig. 3a. Significance was determined by paired t-test. **f)** Proliferation phenotypes of SNP in *GNL3* 5’UTR (rs1108842). The A-allele is associated with higher lithium-induced proliferation, and increased *GNL3* expression in response to lithium. **g**,**h)** NPCs treated with LiCl for 48 hrs with a 2 hr. EdU pulse, followed by labeling with anti-GNL3 antibody and DAPI. (**g**) Representative image demonstrates GNL3 is a nuclear protein. (**h**) Quantification of GNL3 puncta in cells in EdU-cells versus EdU+ cells in the presence and absence of LiCl. n=8 wells per condition, nine tiled images per well. Significance was determined by a paired t-test. Scale bar = 10 um.

GNL3, also known as nucleostemin, regulates cell cycle progression and genome stability, and is essential for maintenance of stem cell fate^43,44^. *GNL3* was also identified as a putative BD risk gene in the most recent BD GWAS^4^, and an integrated eQTL analysis of previous BD GWAS^45^. rs11709284 is an eQTL in GTEx for multiple nearby genes (*PBRM1, NT5DC2, GLYCTK, NEK4*), but most significantly for *GNL3* in brain tissues, including frontal cortex and hippocampus^46^ (Fig. 3b,c), and these associations colocalize (Extended Data Fig. 9a-g). The allele associated with increased *GNL3* is also associated with increased lithium-induced proliferation (Fig. 1f). We next assessed allele specific *GNL3* expression in the population of NPCs using SNPs in LD with rs11709284 in the *GNL3* open reading frame, and found that the allele associated with increased *GNL3* expression at baseline was also associated with increased lithium-induced proliferation^30^ (Fig. 3d, Extended Data Fig. 9h). To assess how lithium affects the expression of each *GNL3* allele, we used allele specific qPCR probes targeting a SNP in the 5’UTR of *GNL3* (*r*^*2*^ = 0.68) (Extended Data Fig. 10a). Again, we observed that the allele associated with increased proliferation rates had increased *GNL3* expression in a temporally controlled manner (Fig. 3e,f). Immunolabeling for GNL3 suggested that cells in S-phase have increased nuclear puncta compared to those which did not (Fig. 3g,h). While lithium increased *GNL3* RNA in bulk qPCR measurements (Fig. 3a), we were not able to detect that lithium further increased GNL3 protein levels in individual cells (Fig. 3h). This suggests that lithium is causing more cells to enter the cell cycle, but is not directly affecting GNL3 expression on an individual cell basis. In all, this data suggests that *GNL3* is a lithium and proliferation responsive gene, and that common genetic variants associated with increased *GNL3* expression are also associated with increased lithium induced NPC proliferation.

Next, we sought to determine whether manipulating *GNL3* expression is sufficient to affect lithium responsive NPC proliferation. We targeted the promoter of *GNL3* with dCas9 fused to either a chromatin repressive (KRAB) or chromatin opening (VP64) domain^47,48^, and identified guide RNAs (gRNAs) capable of decreasing or increasing *GNL3* expression (Extended Data Fig. 10a, Fig. 4a). We found these constructs do not detectably affect expression of genes within a 250kb window surrounding the gRNA target sites, except for *PBRM1*, which shares a bidirectional promoter with *GNL3* (Extended Data Fig. 10a-c). Because *PBRM1* was not found to have lithium responsive gene expression, we focus on the effects of modulating *GNL3* at this locus (Extended Data Figure 8c). Interestingly, changing baseline *GNL3* levels did not affect the relative magnitude of lithium’s activation of *GNL3* expression (Extended Data Fig. 10b).

**Figure 4:**
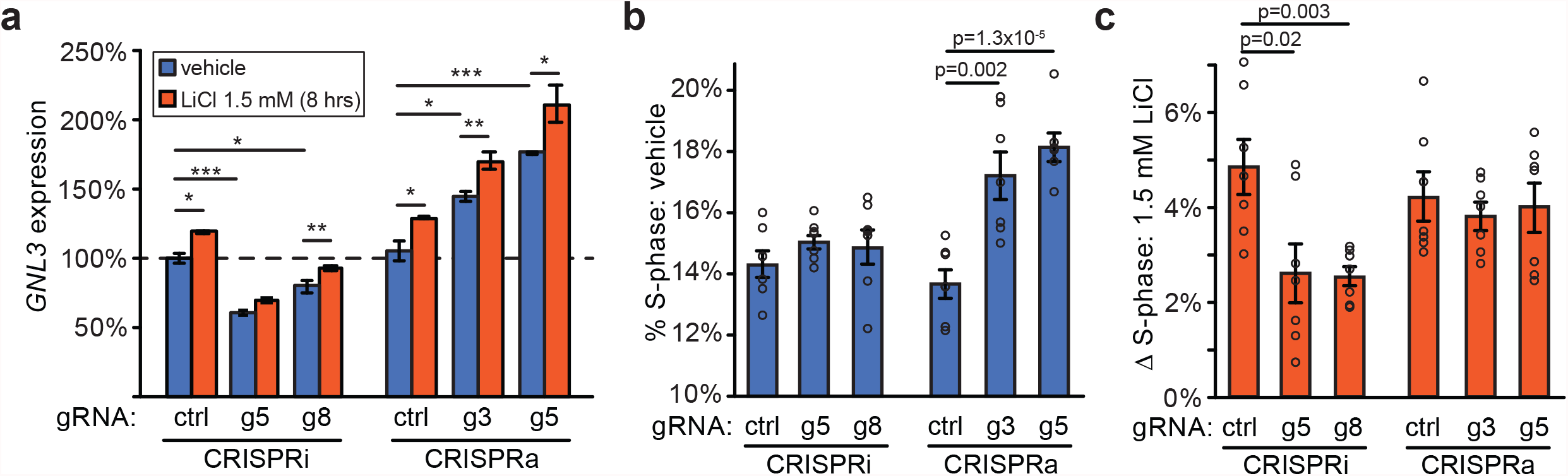
*GNL3* expression is necessary for the full effects of lithium-induced proliferation, and sufficient to induce NPC proliferation. **a)** An NPC donor line heterozygous for index SNP (rs11709284) was transduced with lentivirus carrying the indicated gRNA, and dSpCas9:KRAB (CRISPRi) or dSpCas9:VP64 (CRISPRa). Cells were incubated for four days, followed by an eight hour treatment with 1.5 mM LiCl. *GNL3* expression was analyzed by RT-qPCR. Data normalized to neg. control gRNA (non-targeting, dashed line). n=4 qPCR replicates, significance determined by two sample t-test. * p<0.05, ** p<0.01, *** p<0.001. **b)** Proportion of NPCs in S-phase expressing indicated dSpCas9 and gRNAs at baseline, analyzed by high content imaging, as in Fig. 3g. n=7 wells per condition, nine tiled images per well, significance determined by two sample t-test. **c)** Change in the proportion of NPCs in S-phase in response to 1.5mM LiCl, while expressing indicated dSpCas9 and gRNAs. n=7 wells per condition, significance determined by two sample t-test.

Increasing *GNL3* expression was sufficient to increase proliferation in the absence of lithium, while decreasing *GNL3* expression had no detectable effect on baseline proliferation (Fig. 4b). In contrast, NPCs with decreased *GNL3* expression had 52-54% less lithium-induced proliferation (Fig. 4c). Increasing *GNL3* expression did not further affect lithium’s induction of proliferation (Fig. 4c), which we speculate could be a ceiling effect as the dCas9 VP64 treated NPCs already had ∼4% increased proliferation rate at baseline (Fig. 4b). In all, these experiments indicate that *GNL3* is necessary for lithium’s full proliferative effects in NPCs, and sufficient to induce NPC proliferation.

In summary, we used a GWAS approach in cultured human NPCs to identify a genetic locus impacting lithium-induced proliferation. The association between *GNL3* expression and proliferation was only detectable following lithium treatment (Fig. 1d,f), supporting the hypothesis that certain genetic variants may influence traits only in specific contexts^49^. Our results also demonstrate that individuals with decreased baseline levels of *GNL3* expression in NPCs will be desensitized to lithium’s proliferative effects. These findings suggest two clinically relevant hypotheses. First, levels of *GNL3* expression may modulate the proliferative response to lithium in cortical NPCs present during fetal development, and affect risk for bipolar disorder and schizophrenia later in life. However, given the limited epidemiological evidence for teratogenic effects of lithium on human brain development^18,19,50^, this hypothesis remains difficult to test. Second, assuming that these fetally derived NPCs can model the proliferative activity of adult hippocampal progenitor cells and that proliferation of these cells provide therapeutic benefit as has been shown for antidepressants^26–28^, our results suggest that manipulating *GNL3* expression during lithium treatment may have therapeutic benefits. Future studies can test this hypothesis by directly manipulating levels of *GNL3* in adult hippocampal NPCs during lithium exposure and determining the effects on NPC proliferation, neurogenesis, and rescue of relevant behavioral paradigms. Our study also demonstrates that leveraging the genetic diversity inherent in libraries of primary or iPSC-derived human cell lines allows identification of therapeutically relevant genes and pathways.

## Materials and Methods

### NPC cultures

Primary human NPCs were obtained from fetal brain tissue assumed to be derived from the dorsal telencephalon based on visual inspection at approximately 14-21 gestation weeks, as previously described^30–32^. Tissue was acquired by the UCLA Gene and Cell Therapy Core according to IRB regulations. After single cell dissociation, cells were initially cultured as neurospheres before plating on fibronectin (Sigma, Cat#:F1141) and polyornithine (Sigma, Cat#: P3655) coated plates, where they were passaged 2-3 times, cryopreserved, and transferred to UNC Chapel Hill. NPC media: Neurobasal A (Life Technologies, 10888-022) supplemented with 100 μg ml^−1^ primocin (Invivogen, ant-pm-2), 10% BIT 9500 (Stemcell Technologies, 09500), 1% glutamax (100×; Life Technologies, 35050061), 1 μg ml^−1^ heparin (Sigma-Aldrich, H3393-10KU), 20 μg ml^−1^ EGF/FGF (Life Technologies, PHG0313/PHG0023), 2 ng ml^−1^ LIF (Life Technologies, PHC9481) and 20 ng ml^−1^ PDGF (Life Technologies, PHG1034). We followed previously established protocols to maintain NPCs as proliferating neural progenitors and inhibit differentiation into neurons^31^. No mycoplasma contamination was detected during regular pre-assay screens of cell culture media (ATCC 30-1012K).

### LiCl treatments

This describes the general approach for the lithium exposure performed in all experiments. LiCl (Sigma, 203637) was diluted in water to 3M and stored in single use aliquots at -80·C. NPCs were plated at densities dependent on culture plate size, as described below for each experiment. The following day LiCl was diluted in NPC media, without growth factors, to 10X concentration in 10% of the total volume media in each well. Control wells (vehicle) contained a volume of water equal to the amount of 3M LiCl required to obtain 10X concentration.

### RNA extraction and RT-qPCR

200,000 NPCs were plated in each well of a 12 well plate (Fisher Scientific, Cat#: 0720082). LiCl was added at different times, and RNA for all timepoints was collected simultaneously. Each timepoint had paired vehicle/LiCl exposed samples for normalization purposes. RNA was extracted using standard Trizol extraction method (Ambion). cDNA was synthesized from 100-200 ng RNA using VILO SSIV Master Mix with ezDNAse (ThermoFisher). qPCR was performed with SSO Advanced SYBR Green MasterMix (BioRad). n=4 qPCR replicates were performed for all experiments. Relative gene expression was determined using the ΔΔCt method, normalized to *EIF4A2*. Each LiCl condition was normalized to its respective vehicle time point. Statistical tests for timepoint experiments use the paired Student’s t-test (Fig. 3a,e, Extended Data Figs. 1b,d,e, 8c,d), all other qPCR experiments use the two-sample Student’s t-test (Fig. 4a, Extended Data Figs.1f-j, 10c). qPCR primers are listed in Supplemental Table 3.

### Lentivirus production

Lentivirus was produced in HEK293T cells using the third-generation packaging plasmids (Addgene #12260, #12259). Multiple densities (200,000-600,000 cells) of HEK293T cells were plated in 12 well plates in 1 mL media. 24 hours later the cell density at ∼90% confluency was transfected with 1200 ng psPAX2, 800 ng pMD2.G, and 1600 ng of each lentiviral plasmid, using 8 uL FuGene6 (Promega). Cells were incubated for 24 hours, followed by a 50% media change in the morning, and a 1 mL media addition in the afternoon. 24 hours later supernatant was collected, filtered using 0.45 μm filters, and stored in single use aliquots at -80·C.

### Wnt signaling luciferase assays

10,000 NPCs per well were plated in 96 well plates (Corning, Cat# 07-200-91), and 24 hours later were transduced with 10 uL lentivirus carrying BAR:luciferase lentivirus, and 2 uL Tk:*Renilla* lentivirus. Cells were incubated for two days, treated with LiCl as above, and incubated for 48 hours. Cell lysate was used in dual luciferase assays using the Dual-Glo luciferase system (Promega), measured on the GloMax Discover plate reader (Promega). BAR:Luciferase and Tk:*Renilla* plasmids were kind gifts from the lab of Ben Major^51^. Statistical significance was determined using paired Student’s t-test.

### NPC line selection for GWAS and QC

Cell type heterogeneity in the NPC library could introduce phenotypic variation based on cell type, which could be a confounding factor in genetic associations. Proliferation assays were performed on 94 NPC donor lines, but we excluded NPC donors with outlier gene expression patterns likely resulting from cell-type heterogeneity due to errors in dissection. Principal component analysis was performed on previously acquired^30^ baseline expression of the 500 highest variance transcripts followed by k-means clustering (k=2) (Extended Data Fig. 3a-c). This analysis identified NPC donor lines with relatively low expression of the canonical NPC marker *PAX6*, and relatively high levels of *NKX2*.*1* and *VAX1*, markers of ventral inhibitory NPCs^52,53^. To assess sample swaps or mixing between NPC lines we used verifyBamID^54^, which flagged an additional 4 lines with FREEMIX or CHIPMIX scores greater than 0.04. This resulted in NPCs from 80 distinct donors that were carried forward into association analyses.

### High-throughput proliferation assays

We thawed cryopreserved NPCs in batches of ∼8-10 NPC donor lines, which were pseudorandomized for biological and technical variables (sex, gestation week, passage). We included at least one technical replicate (distinct cryovials of the same NPC donor line thawed and assayed multiple times) in each batch to assess technical reproducibility (29 unique NPC donor lines replicated in duplicate, Extended Data Fig. 3h). Each batch was cultured with weekly passaging for two weeks to allow recovery from thawing. On week three, cells were lifted using 0.05% Trypsin (Gibco), and 12,500 cells per well were plated in each well of 96 well plates (Corning, Cat. #: 3610). 24 hours later, LiCl (0.75, 1.5, or 3 uM) or vehicle (water) was added as described above. After 46 hours, cells were treated with 10 uM EdU with a 10% media addition, and incubated for 2 hours. Cells were then lifted off the plate using 0.05% trypsin (Gibco), and transferred to a u-bottom 96-well plate. Cells were fixed with 4% PFA in PBS for 10 minutes. EdU labeling was performed using the Click-iT EdU Cell Proliferation Kit (ThermoFisher, C10337) per manufacturer’s protocols. Total DNA content was labeled with the FxCycle Far Red dye (ThermoFisher, F10347). Cell suspensions were quantified using the Attune NxT 96-well Flow Cytometer. For each LiCl concentration, 4 wells per NPC donor line were quantified. For vehicle (water), 8 wells per NPC donor line were acquired. Replicates were spatially distributed across each plate to avoid positional bias. All liquid handling steps, including the additions of LiCl, EdU, trypsin, fixation, and the Click-iT reaction, were performed using the Tecan Evo liquid handling robot to reduce handling variability.

### Analysis of flow cytometry data

FCS output files from the Attune NxT were initially analyzed using FlowJo, using SSC to retain only singlets (Extended Data Fig. 2a,b). To perform automated gating of stages of the cell cycle, we combined technical replicates into a single FCS file, and bounds were drawn for G0/G1, S-phase, and G2/M using the automated gating software FlowDensity^55^, with three non-elliptical gates (Extended Data Fig. 2c). These gates were then applied to each well independently. Technical replicates were averaged to obtain the percentage of cells in S-phase for each NPC donor line in each condition. Wells which failed for technical reasons (such as a bubble in the flow cytometer or no EdU labeling due to a failed Click-iT reaction), were identified by calculating the coefficient of variation (CV), followed by manual inspection of FCS files for all conditions where CV > 0.1. This filtering removed ∼0.5% of wells. For donor lines which were assayed in multiple rounds, we averaged all replicates together for genetic associations.

### Genome-wide association

The percentage of cells in S-phase for each experimental condition for each NPC donor line was calculated using the average across all wells. ΔS-phase was calculated by subtracting the percentage of cells in S-phase in vehicle condition from the percentage of cells in S-phase for each concentration of LiCl. We used a linear mixed effects model to conduct genetic association tests implemented with EMMAX software^37^. In this model, we included an *m x m* Balding Nichols kinship matrix inferred from NPC genotypes (*K*)^56^ as a random effect variable to control for effects of cryptic relatedness, and 10 multidimensional scaling (MDS) genotype components as covariates to mitigate confounding effects of population structure. While we did not find that sex or gestation week correlated with proliferation phenotypes (Extended Data Figs. 3j-l), we included sex and gestation week as standard technical covariates in the model.

Study-wide significance threshold (as in Nyholt, *et al*^*38*^) was used because we ran four phenotypes (S-phase: vehicle, ΔS-phase 0.75 mM, 1.5 mM, 3 mM LiCl). In brief, we first calculated the total number of independent tests across these GWAS using summary statistics (determined to be 2.996 independent tests), and then adjusted the standard genome-wide significance threshold (*γ*= 5×10^−8^) by the effective number of independent tests (*n*), to derive a study-wide significance threshold of *α* = 1− (1− *γ*)^1/*n*^ = 1.67×10^−8^.

### GWAS colocalization

We evaluated numerous GWAS summary statistics (Supplemental table 2) for colocalization with the study-wide significant locus on chromosome 3. We first filtered downloaded GWAS summary statistics for SNPs with genome-wide significant p-values (p < 5×10^−8^) within 1Mb upstream or downstream from rs11709284. Next, we calculated LD between these SNPs and rs11709284 using either our population of NPC donors or 1000 Genomes Project Europeans (EUR)^57^. We required variants to exceed *r*^*2*^ > 0.6 to be considered a colocalization candidate, and then used conditional analysis to provide evidence for colocalization^58^. We considered GWAS signals to be colocalized when the proliferation associated p-value no longer reached nominal significance (p > 1 × 10^−5^) after conditioning on the GWAS index SNPs.

### Cross-trait enrichment analysis

GWAS summary statistics (Supplemental Table 2) were filtered for variants also associated with proliferation phenotypes from this study under increasingly stringent p value thresholds (p < 1×10^−3^, 1×10^−4^, 1×10^−5^, 1×10^−6^, 1×10^−7^). Observed distributions of filtered association p values from the various GWAS were compared to expectations under a null hypothesis where no genetic effect on the phenotype exists using quantile-quantile plots. Genomic inflation factor^59^, *λ*_*GC*_, was calculated from filtered p values to quantify enrichment of associations across traits.

### Allele-specific gene expression via RNAseq and qPCR

ASE analysis was performed as described in Aygün et al. 2021^30^ using variants supported by at least 10 allele-specific counts in total (at least 2 from either allele) from each of the heterozygous donors in the analysis. 3 NPC donor lines heterozygous for rs1108842 (*r*^*2*^_rs11709284,rs1108842_ = 0.68) were plated in 12 well plates and treated with LiCl as described above. cDNA was generated from 200 ng total RNA using VILO SSIV Mastermix (ThermoFisher). Allele specific expression levels were determined using TaqMan Universal Master Mix II (ThermoFisher), and TaqMan genotyping probes for rs1108842 (Applied Biosystems). Quantification of allele specific expression was calculated by assessing the difference between Ct values for each allele within each PCR well at ΔRn=0.5, as previously described^60^. Experiments were performed in three distinct NPC donors, n=4 technical replicates per NPC donor line. Each LiCl time point was normalized to its respective vehicle time point, and then further normalized to the 0 hour time point. Statistical significance was determined using paired Student’s t-test.

### CRISPR based modulation of gene expression

gRNAs were selected using the GPP sgRNA Designer (Broad Institute) and cloned into CRISPRi (pLV hU6-sgRNA hUbC-dCas9-KRAB-T2a-GFP; Addgene # 71237) or CRISPRa (pLV hU6-gRNA (anti-sense) hUbC-VP64-dCas9-VP64-T2A-GFP; Addgene #66707) plasmids, which were kind gifts from the lab of Charles Gersbach. Lentivirus was prepared as above. For experiments in Fig. 4a an NPC donor line heterozygous for rs11709284 was plated at 200,000 cells per well in 12 well plates. 75 uL of each viral prep was delivered to each well. Cells were incubated for 96 hours. RNA extraction, cDNA synthesis, and qPCR were performed as described above. Consistency of lentivirus infection and dCas9 expression was verified using qPCR for dCas9. The negative control gRNA is a random sequence that does not align to the human genome.

### Quantifying proliferation via EdU incorporation and high content imaging

For experiments in Fig. 3g,h, a randomly selected NPC line was plated at 10,000 cells per well in black walled 96 well plates (Corning, Cat#: 3603). 24 hours later LiCl was added to the plate as described above and incubated for 48 hours, with an EdU pulse during the last two hours of incubation. For experiments in Fig. 4b,c, a randomly selected NPC donor line was plated at 200,000 cells per well in 12 well plates. 75 uL of each CRISPRi/a viral prep was delivered to each well. Cells were incubated for 96 hours. Cells were then lifted off the plate using 0.05% trypsin, counted, and plated in black walled 96-well plates at 10,000 cells per well. 24 hours later cells were treated with LiCl at the indicated concentrations, and incubated for 48 hours with an EdU pulse during the last two hours of incubation. Cells were fixed on the plate with 4% PFA in PBS, and Click-iT EdU labeling was performed per the manufacturer’s protocol (ThermoFisher: C10337). Cells were additionally labeled with anti-GNL3 antibody (ThermoFisher: AB_2532414, clone:3H20L2, 1:250 dilution), anti-GFP antibody to label Cas9+ cells (AbCam: ab5450, 1:1000 dilution) and DAPI (1:4000). Images were acquired at 20x using the GE IN CELL Analyzer 2200 high-content imager. Images were analyzed using a custom CellProfiler pipeline. We isolated Cas9+/DAPI+ positive cells, and quantified the number of GNL3 puncta per nucleus (puncta defined as nuclear object 3-10 pixels in diameter), and percentage of Cas9+/EdU+ cells using n=7-8 wells per condition, with 9 images captured per well. Cell counts of each image were averaged for each well, and wells were averaged across conditions. Significance was determined by a paired t-test, paired by well.

## Figure legends

**Extended Data Figure 1:**
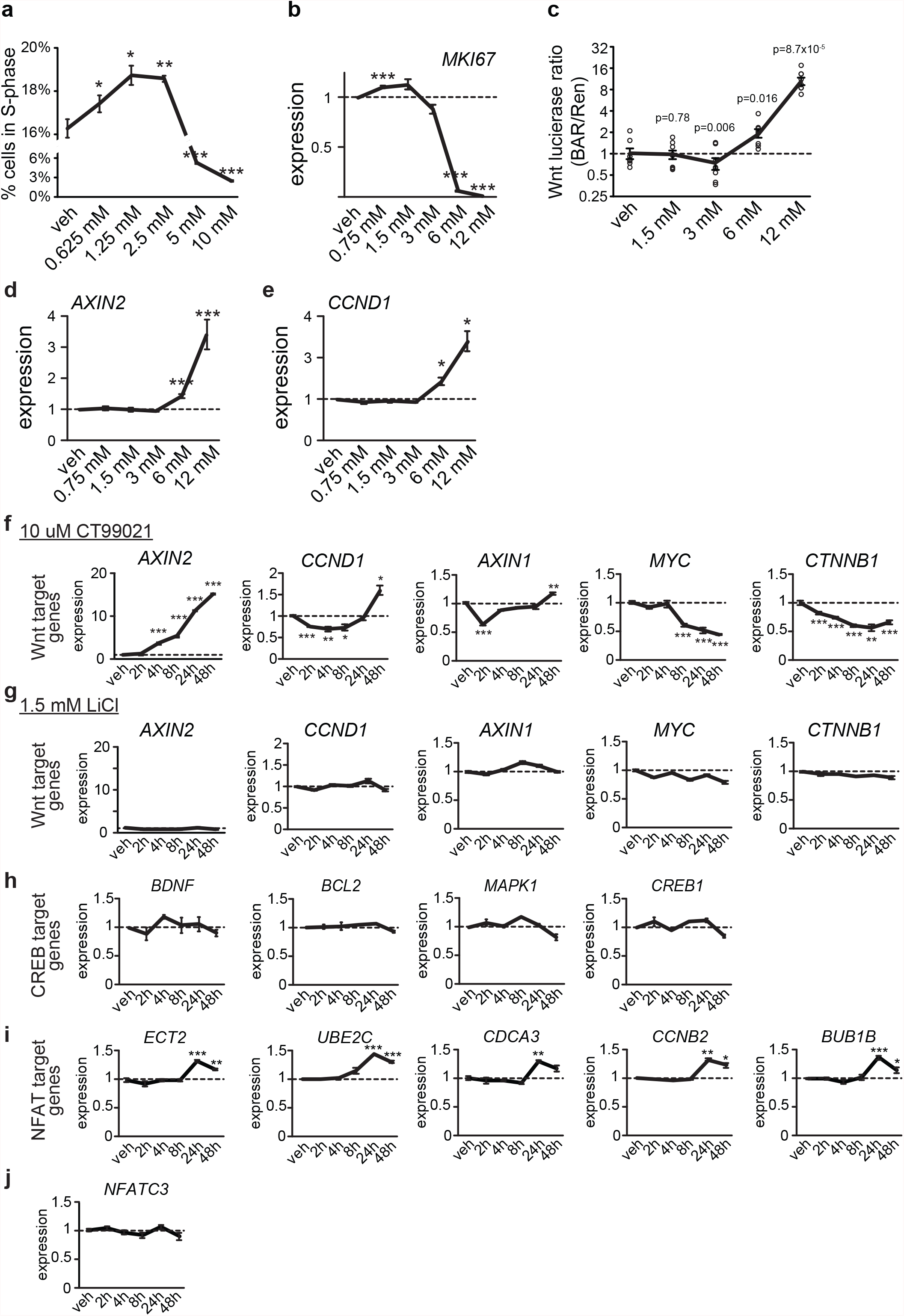
Effects of lithium on NPCs. **a)** NPCs from one donor were exposed to LiCl for 48 hours, with an EdU pulse during the last two hours to label cells in S-phase. Cells were then lifted off the plate, fixed, labeled with a DNA content dye, and analyzed by flow cytometry. n=4 wells per concentration. Significance was determined by two sample t-test. All error bars are standard error of the mean. * p<0.05, ** p<0.01, *** p<0.001. **b)** NPCs from three donors were exposed to LiCl for 48 hours at the indicated concentrations, followed by RNA extraction and RT-qPCR using primers for the proliferation marker gene *MKI67*. Data was normalized to the housekeeping gene *EIF4A2*. n=4 per wells per concentration. Significance was determined by paired t-test. **c)** NPCs from 8 distinct donors were transduced with lentivirus carrying Firefly luciferase under control of the TCF/LEF sensitive promoter to measure Wnt signaling activity and a constitutively active Renilla luciferase to control for level of transduction. 48 hours later cells were treated with the indicated concentration of LiCl, followed by a 48 hour incubation. Cell lysate was then subjected to a dual luciferase assay. Firefly luciferase was normalized Renilla luciferase in each well. Data were normalized to vehicle. n=4 technical replicates per condition, significance at each concentration compared to vehicle was determined by paired t-test. **d-e)** NPCs from three donors treated with LiCl for 48 hours at the indicated concentrations, followed by RNA extraction and RT-qPCR using primers for the indicated Wnt pathway associated genes. Data were normalized to the housekeeping gene *EIF4A2*. n=4 per condition, significance determined by paired t-test. **f-j)** Gene expression in one NPC line across multiple timepoints following treatment with the highly selective Wnt activator CT99021 (**f**), or 1.5 mM LiCl (**g-j**). **f**,**g**: target genes of Wnt signaling pathway, **h**: target genes of the CREB signaling pathway, **i**: target genes of the NFAT signaling pathway^35^, **j**: *NFATC3* transcription factor. Data normalized to the vehicle sample that was extracted at the same time (dashed line). n=4 wells per concentration. Significance was determined by two sample t-test.

**Extended Data Figure 2:**
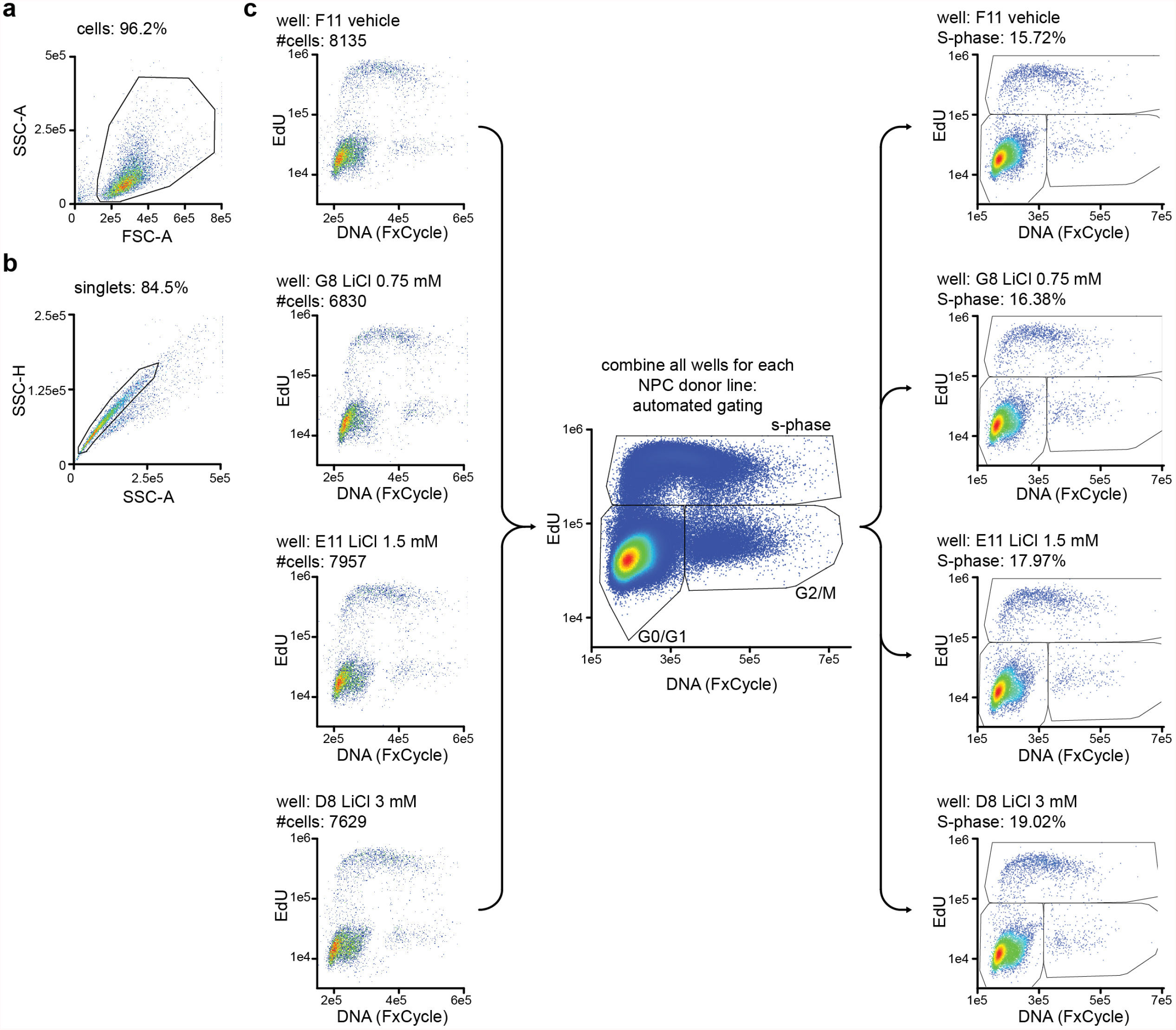
Flow cytometry gating strategy. **a)** Density plots of a well from a representative NPC donor line. Forward scatter area (FSC-A) and side scatter area (SSC-A) were used to gate around cells and remove debris from analysis. This gate was manually drawn for a vehicle well, and applied to all wells corresponding to that NPC donor line. The percentage refers to the percent of events within the gate. **b)** SSC height and SSC-A were used to isolate singlets. **c)** Representative density plots from single cells isolated from **b**, for arbitrarily selected wells treated with each experimental condition. All wells from each NPC donor line were combined into a single FCS file, and FlowDensity^55^ was used to automatically draw boundaries between cells in each phase of the cell cycle. These gates were then applied to each well individually to quantify the percentage of cells in S-phase.

**Extended data Figure 3:**
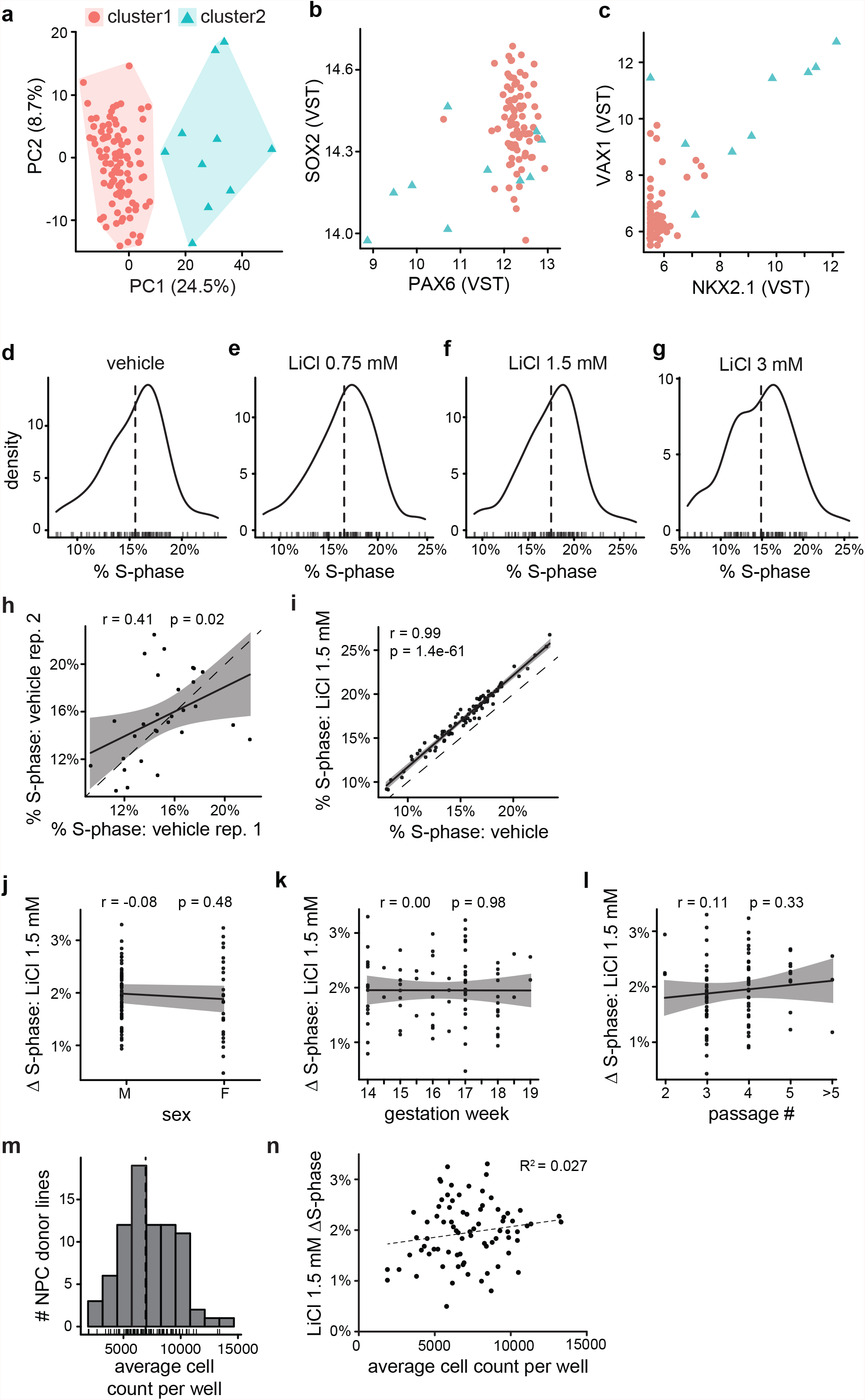
Proliferation phenotype quality control. **a)** To identify transcriptional heterogeneity in the library of NPCs we performed principal component analysis (PCA) of baseline gene expression^30^ of the 500 highest variance genes in NPCs from 94 distinct NPC donor lines. Each donor line is a single point plotted in PC space. Clusters were identified by k-means clustering (k=2). Genetic association was performed in NPCs from cluster 1. **b)** Scatterplot of VST-normalized *SOX2* and *PAX6* expression across NPCs in each cluster suggests that cluster 2 are outliers for expression of canonical NPC marker genes *PAX6* and *SOX2*. **c)** Scatterplot of VST-normalized *VAX1* and *NKX2*.*1*, which are markers of cells derived from the medial ganglionic eminence (MGE) in the ventral telencephalon. **d-g)** Distribution of proliferation rates across NPCs from 80 distinct donor lines at the indicated LiCl concentrations. Approximate normality suggests proliferation phenotypes are amenable to linear modeling in genetic association tests. **h)** Correlation between technical replicates, defined as NPCs from the same donor line thawed and assayed in two different batches. Dashed line, y=x. **i)** Correlation between proliferation in vehicle and 1.5 mM LiCl conditions. Dashed line, y=x. **j-n)** Correlation between proliferation rate and technical variables: sex (**j**), gestation week (**k**), passage number (**l**), and average number of cells in each well (**m**,**n**).

**Extended Data Figure 4:**
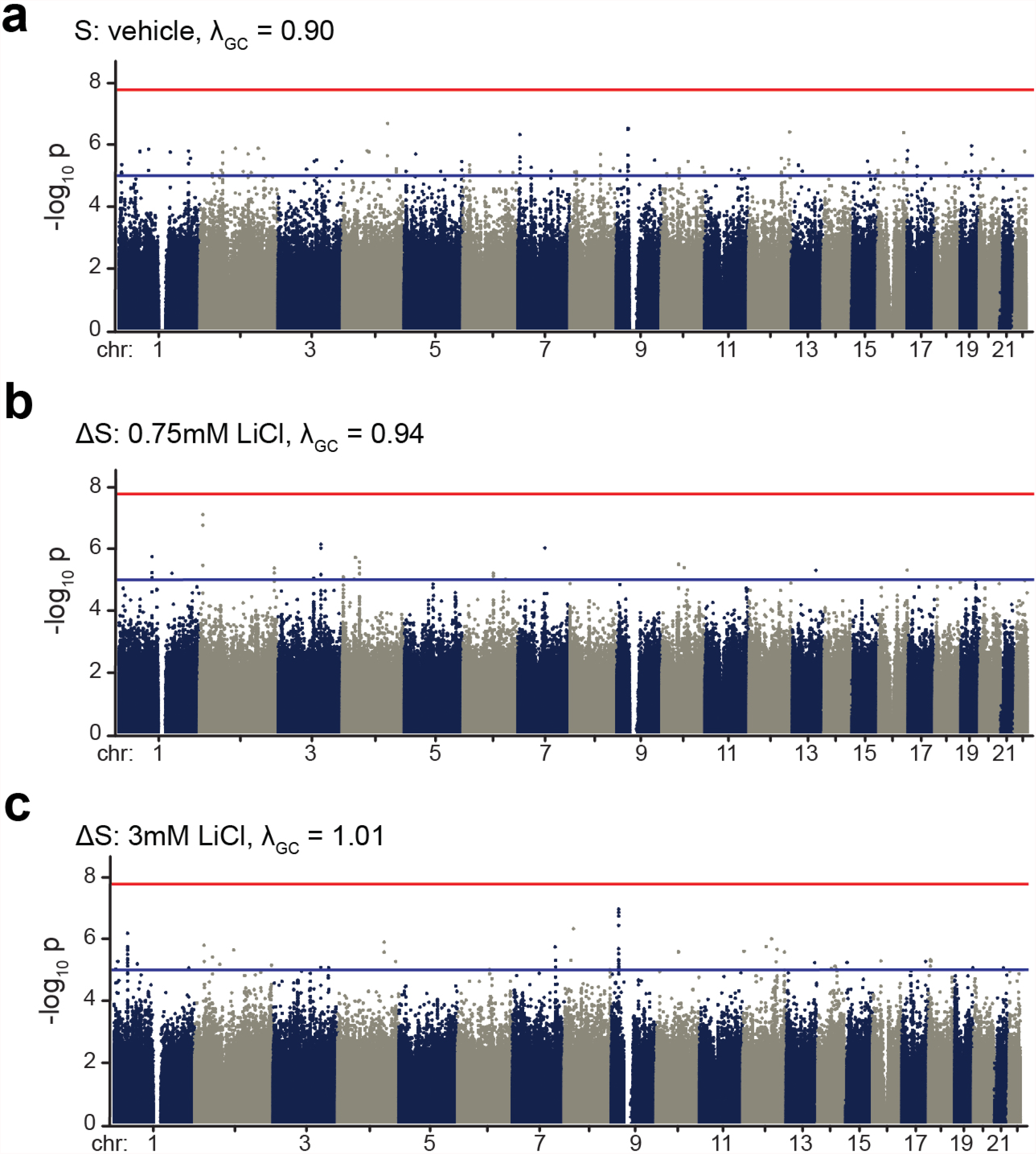
Lithium induced proliferation GWAS results. **a-c)** Manhattan plots of GWAS results from vehicle S-phase (**a**), 0.75mM LiCl ΔS-phase (**b**), and 3mM LiCl ΔS-phase (**c**). Red line denotes study-wide significance level (P < 1.67 × 10^−8^); blue line denotes nominal significance level (p < 1 × 10^−5^)

**Extended Data Figure 5:**
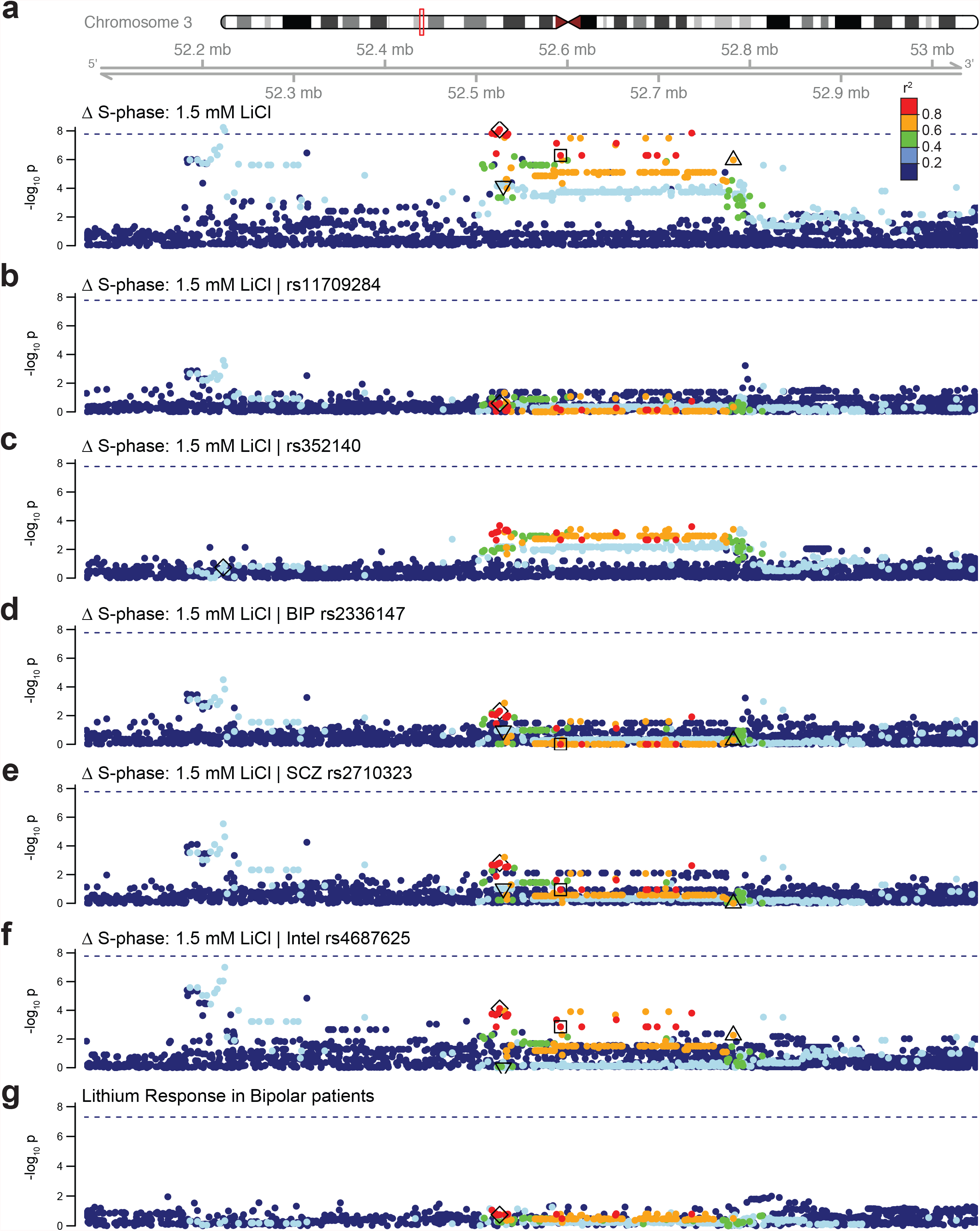
GWAS results at chr3p21.1 - conditional tests and lithium response. **a)** Locus zoom of the study wide significant locus on chr3p21.1, associated with 1.5 mM LiCl ΔS-phase phenotype (Same data as Fig 2a). ◊: SNP from this study (rs11709284). Dashed line denotes the study-wide significance threshold (p < 1.67×10^−8^) for all plots. For all plots, variants are colored by LD relative to rs11709284 in NPCs used for this study. **b)** Locus zoom of 1.5 mM LiCl ΔS-phase phenotype while conditioning on rs11709284 (**b**) and rs352140 (**c**). **d-f)**Locus zoom of 1.5 mM LiCl ΔS-phase phenotype while conditioning on indicated SNPs from colocalized traits: BD^4^ (**d**), schizophrenia^41^ (**e**), and intelligence^42^ (**f**). SNP annotations: ◻: index SNP for BD (rs2336147). △: index SNP for schizophrenia (rs2710323). ▽: index SNP for intelligence (rs4687625). **g)** Locus zoom for GWAS summary stats from the ConLiGen GWAS on lithium response in individuals with Bipolar disorder (Continuous variable, all populations)^9^.

**Extended Data Figure 6:**
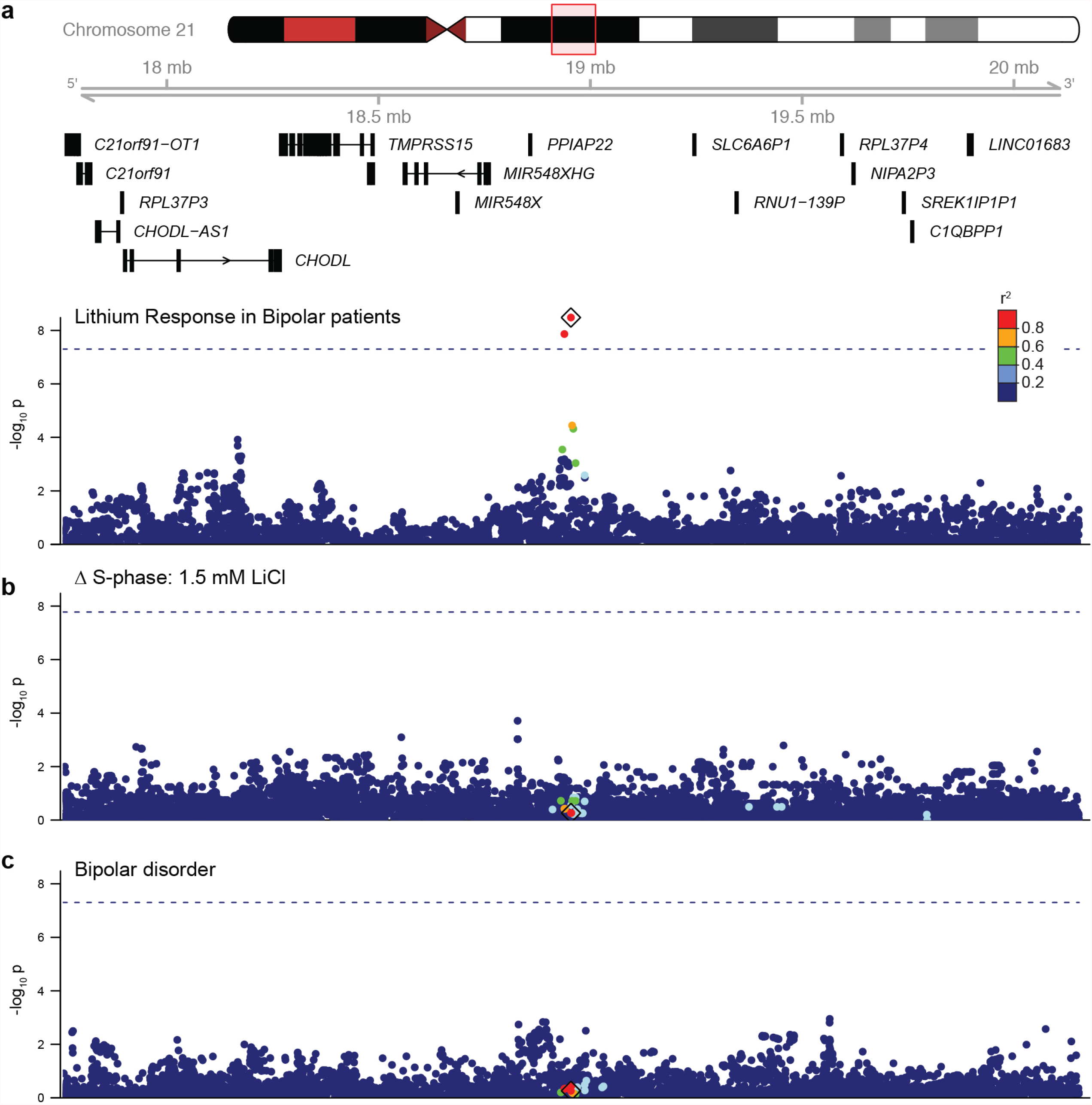
A locus associated with lithium response in individuals with bipolar disorder does not colocalize with lithium-induced proliferation or Bipolar disorder risk. **a)** Locus zoom plot of Lithium response in bipolar patients phenotype at chr21q21.1 (Continuous variable, all populations)^9^. ◊: index SNP (rs74795342). Each SNP is colored by LD (*r*^*2*^) to rs74795342 in 1000 Genomes ALL reference panel. Dashed line denotes genome-wide significance threshold (p < 5×10^−8^). **b)** Locus zoom plot of 1.5 mM LiCl ΔS-phase phenotype. Dashed line denotes the study-wide significance threshold (p < 1.67×10^−8^). **c)** Locus zoom plot of Bipolar Disorder GWAS results^4^. Dashed line denotes genome-wide significance threshold (p < 5×10^−8^).

**Extended Data Figure 7:**
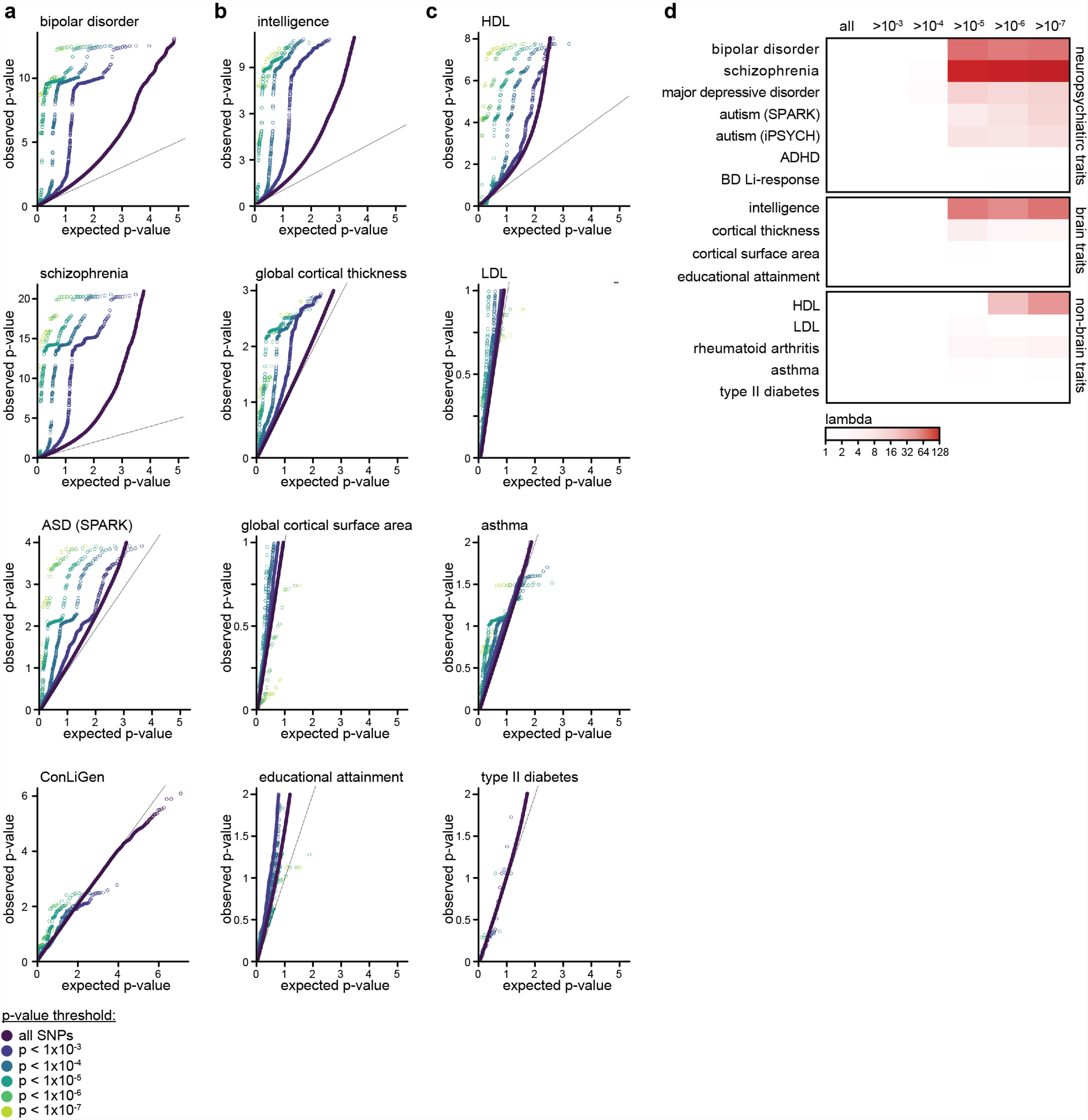
QQs. **a-c)** Quantile-quantile (QQ) plots of association p-values for neuropsychiatric disorder traits **(a)**, brain traits **(b)**, and non-brain traits **(c)** (references for GWAS used found in Supplemental Table 2). Point color denotes SNPs passing increasing filtering strength based on p-value of association to 1.5 mM LiCl ΔS-phase. Each plot depicts the expected distribution of test-statistics on the x-axis vs. the observed test-statistics for the indicated trait on the y-axis. To visualize enrichment of significant GWAS results, y-axes were scaled to the maximum -log_10_(p value) for GWAS sumstats associated with 1.5mM LiCl ΔS-phase phenotype at a threshold of p < 1 × 10^−7^. **d)** Lambda GC^59^ values for each GWAS filtered by ΔS-phase 1.5 mM LiCl GWAS results at increasing p value thresholds. Numbers in parentheses reflect total GWAS sample size for each trait. *sample size for ASD SPARK meta-analysis^62^ includes 6222 case-pseudocontrol pairs.

**Extended Data Figure 8:**
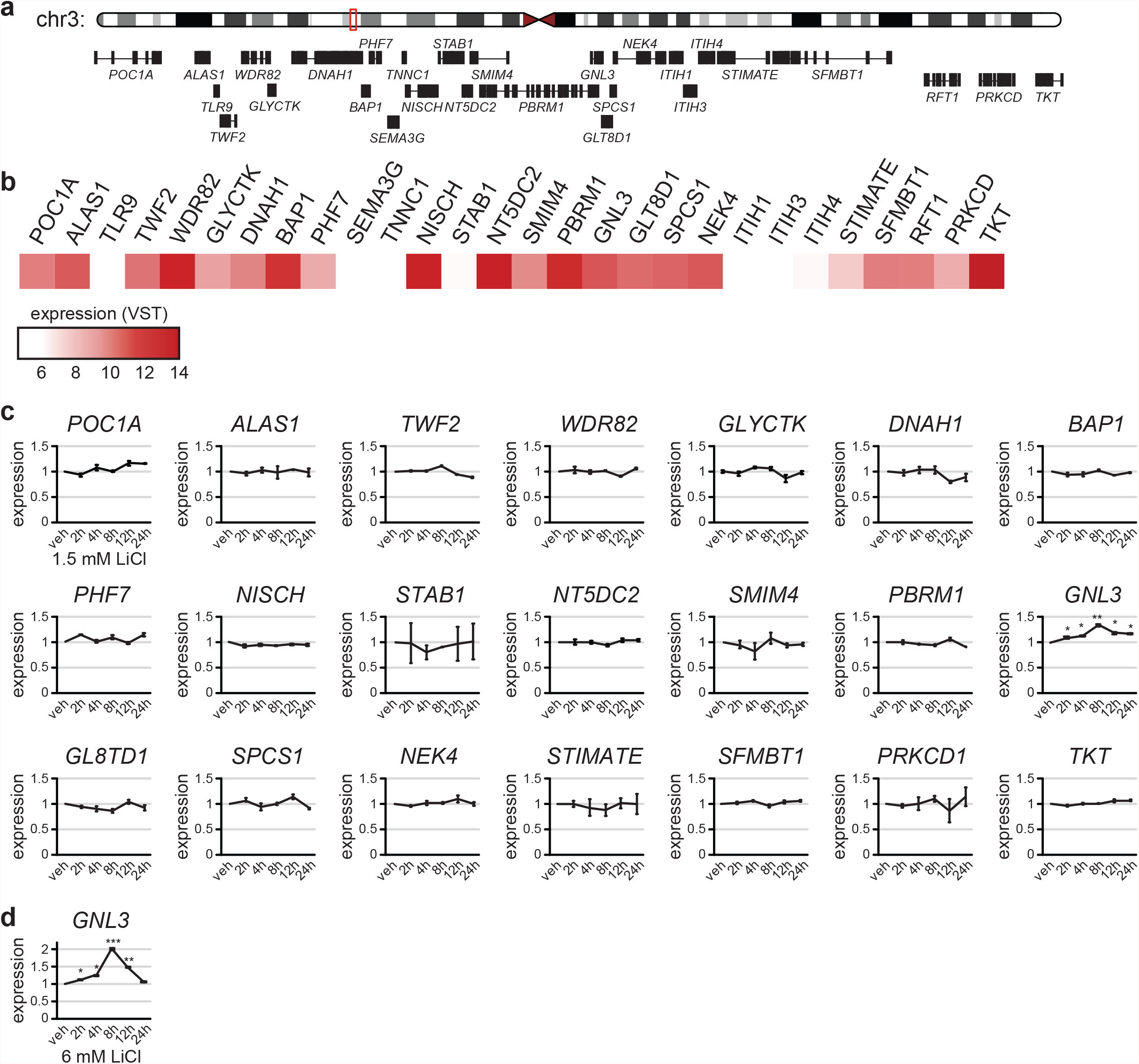
Lithium induced gene expression of genes at the associated Chr3 locus. **a)** 1.2 Mb window surrounding the Chr3 locus, with gene models for all protein coding genes. **b)** Expression levels (VST) of all genes in the window from ^30^. **c)** RT-qPCR from one NPC donor line of all genes with detectable expression at baseline, in response to 1.5 mM LiCl at various time points. *GNL3* is the only gene whose expression changes at multiple timepoints. Data analyzed as in Extended Data Fig. 1d-h. n=4 technical qPCR replicates. Statistical significance assessed by paired Student’s t-test * p<0.05, ** p<0.01, *** p<0.001. **d)** *GNL3* expression in response to 6 mM LiCl.

**Extended Data Figure 9:**
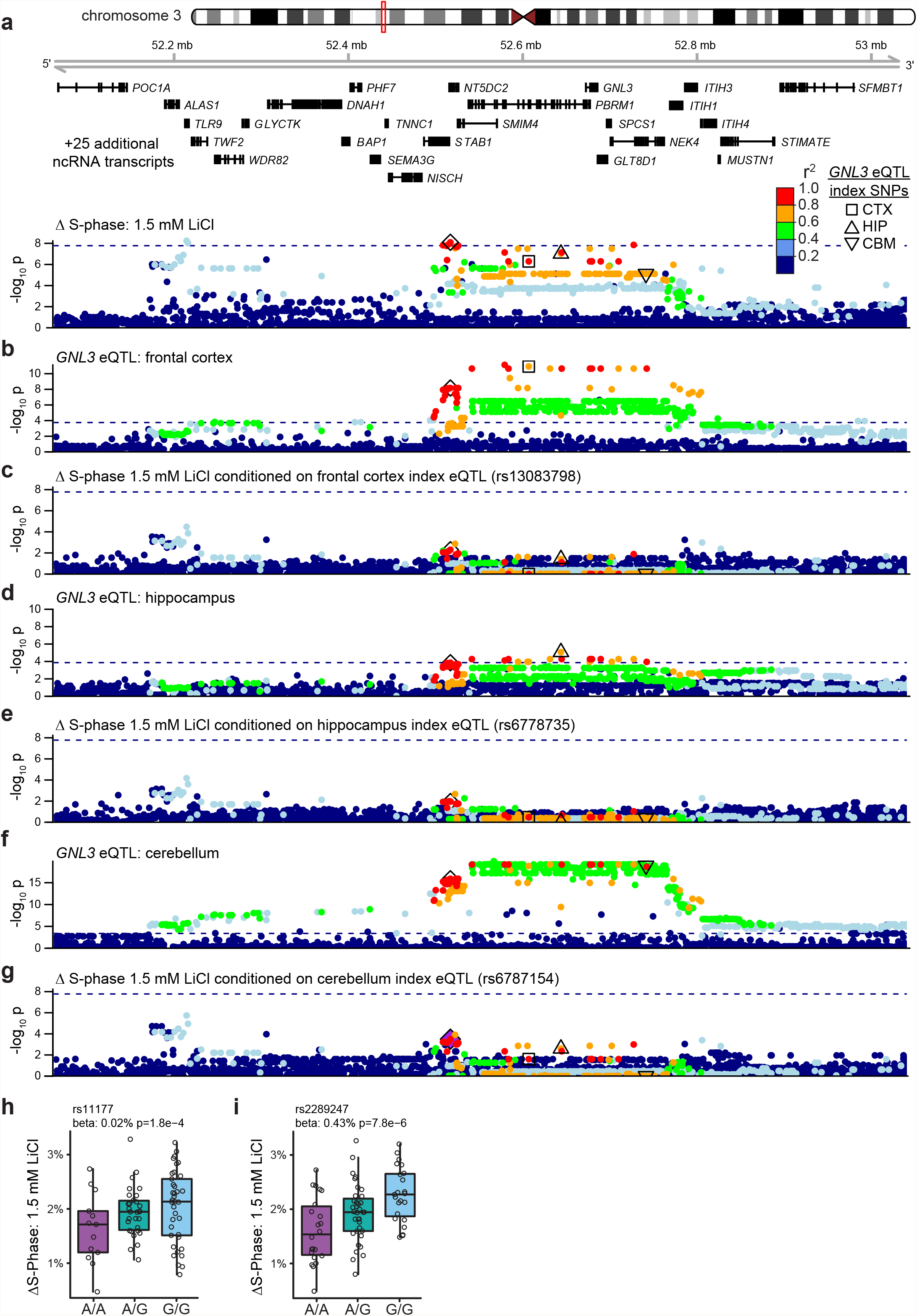
Colocalizations with GNL3 eQTL. **a)** Locus zoom of the study wide significant locus associated with 1.5 mM LiCl ΔS-phase phenotype (Same data as Fig 2a/b). ◊: SNP of interest from this study (rs11709284). ◻: GTEx index eQTL SNP from Frontal Cortex (BA9) (rs13083798). △: GTEx index eQTL SNP from hippocampus (rs6778735). ▽: GTEx index eQTL SNP for cerebellum (rs6787154). Variants are colored by LD relative to rs11709284 in NPCs used for this study. Dashed line denotes the study-wide significance threshold (p < 1.67×10^−8^). **b)** GTEx eQTL for GNL3 (EUR) from Frontal Cortex (BA9) brain tissue. Dashed line denotes the eQTL significance threshold (p < 0.000176, FDR < 0.05). Variants are colored by LD relative to rs11709284 in the 1000 Genomes Project reference panel (EUR). **c)** GWAS results as in (a), conditioned on GTEx eQTL index SNP from Frontal Cortex (BA9) brain tissue (rs13083798). Variants are colored by LD relative to rs11709284 in NPCs used for this study and annotated as in panel (a). Dashed line denotes the study-wide significance threshold (p < 1.67×10^−8^). **d)** GTEx eQTL for GNL3 (EUR) from hippocampus. **e)** GWAS results as in (a), conditioned on GTEx eQTL (EUR) from hippocampus brain tissue (rs6778735). **f)** GTEx eQTL for GNL3 (EUR) from cerebellum brain tissue. **g)** GWAS results as in (a), conditioned on GTEx eQTL (EUR) from cerebellum brain tissue (rs6787154). **h**,**i)** Boxplots showing proliferation across genotypes at SNP rs11177 (**h**) and rs2289247 (**i**). Related to Fig. 3d.

**Extended Data Figure 10:**
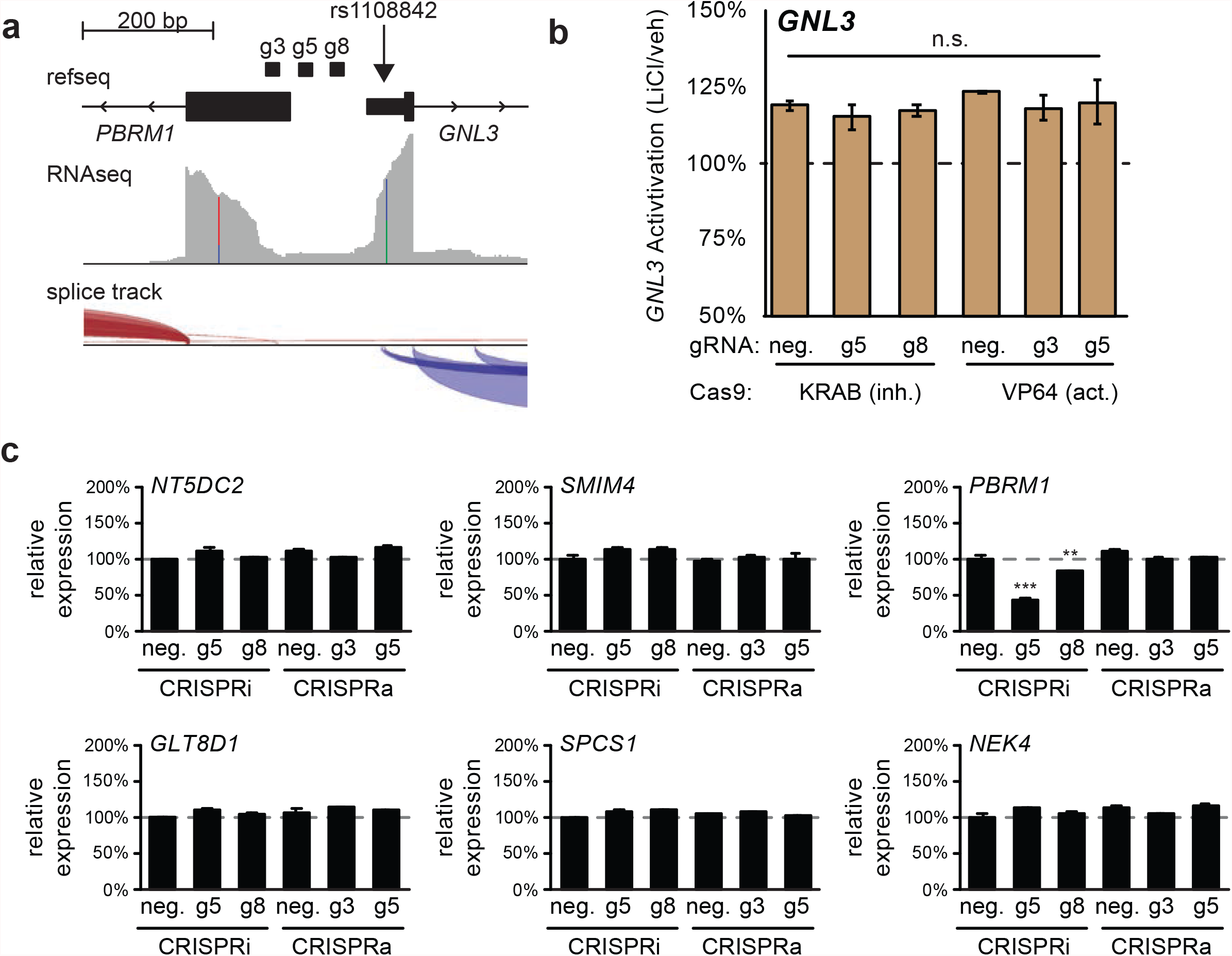
CRISPRi/a targeting *GNL3* in NPCs. **a)** gRNA locations targeting the *GNL3* promoter. rs1108842 is the SNP in the *GNL3* 5’UTR used to assess allele specific expression in Fig. 3e. **b)** *GNL3* expression in an NPC donor line transduced with the indicated lentivirus, and treated with 1.5 mM LiCl. Data normalized to vehicle condition, related to Fig. 4a. Results suggest that regardless of baseline expression, lithium increases *GNL3* expression to uniform levels. n=4 technical qPCR replicates, significance determined by two sample Student’s t-test. **c)** RT-qPCR for the six nearest genes to gRNA target sites other than GNL3, to test for off target activity. Expression normalized to *EIF4A2*. n=4 technical qPCR replicates, significance determined by two sample Student’s t-test. ** p<0.01, *** p<0.001.

**Supplemental Table 1: Nominally significant clumped GWAS results**

Nominally significant (p < 1×10^−5^) GWAS loci from this study clumped using PLINK with 250kb windows and LD *r*^*2*^ > 0.2. Each row reports the most significantly associated variant at each locus. Columns: Trait - associated proliferation phenotype, SNP - variant ID, rsid - variant rsID, CHR - chromosome, BP - chromosomal base-pair coordinates (hg38), P - association p-value, SEbeta - standard error adjusted effect size, TOTAL - number of SNPs in locus clumped with *r*^*2*^ > 0.2, A1 - non-effect (typically minor) allele, A2 - effect allele, freqA1_WntI_n80 - reference allele (A1) frequency within donor-derived NPC lines in this study, freqA1_1kgALL - reference allele (A1) frequency in all subjects from the 1000 Genomes Project reference panel, freqA1_1kgEUR - reference (A1) allele frequency in European subjects from the 1000 Genomes Project reference panel.

**Supplemental Table 2: GWAS summary statistics used for colocalization analysis and cross-trait enrichments**

Columns: trait - description of GWAS trait, PMID/ref - PubMed ID or link to reference, Year - year of study, sample size - total number of subjects in study, nCase - number of cases in study, nControl - number of Control subjects in study, note - additional information about subjects and sample sizes

**Supplemental Table 3: qPCR primers and gRNA sequences used in this study**

Columns: Name - Forward, “F” and Reverse “R” qPCR primers labeled by target gene (rows 2-83), or guide RNAs targeting the GNL3 transcriptional start site (rows 84-86), Sequence -nucleotide sequence of primer or gRNA

## Data and Code Availability

The genotyping data used in this manuscript can be accessed via dbGaP (phs002493.v1.p1 and phs001958.v1.p1). Phenotypic data, code, and GWAS summary statistics available at https://bitbucket.org/steinlabunc/wolterle2022/src/master/

## Author Contributions

JMW, BDL, and JLS designed the experiment and prepared the manuscript. JMW, BDL, and KC performed the proliferation assays in NPCs. JMW performed functional validation and CRISPR based assays. BDL, NM, MJL, NA, DL performed bioinformatic analyses. JLS, JP, and MJZ secured funding to support the work. JLS oversaw the work.

## Acknowledgements

This work was supported by the Foundation of Hope (to JP), the NIH (R01MH120125, R01MH121433, R01MH118349 to JLS; T32HD040127 to JW, T32GM67553-15 to BDL), and the Pfizer-NCBiotech Distinguished Postdoctoral Fellowship in Gene Therapy (to JW). Assistance for this project was provided by the UNC Intellectual and Developmental Disabilities Research Center (NICHD; P50 HD103573; PI: Joseph Piven). We thank Tammy Havener and the UNC Catalyst for Rare Diseases for use of their high-throughput screening equipment. We thank Sergi Papiol, Thomas Schulze, and Francis McMahon of ConLiGen for sharing BD Li-responder GWAS summary statistics ^9^. We thank Janet Dow of the UNC Flow Cytometry Core Facility, which is supported in part by The National Cancer Institute (P30CA016086), awarded to the UNC Lineberger Comprehensive Cancer Center. The UNC Neuroscience microscopy core is supported by the NICHD (U54HD079124).

## References

1. Geddes, J. R. & Miklowitz, D. J. Treatment of Bipolar disorder. Lancet 381, 1672–1682 (2013).

2. Geddes, J. R., Burgess, S., Hawton, K., Jamison, K. & Goodwin, G. M. Long-term lithium therapy for Bipolar disorder: systematic review and meta-analysis of randomized controlled trials. Am. J. Psychiatry 161, 217–222 (2004).

3. Malhi, G. S., Gessler, D. & Outhred, T. The use of lithium for the treatment of Bipolar disorder: Recommendations from clinical practice guidelines. J. Affect. Disord. 217, 266–280 (2017).

4. Mullins, N. et al. Genome-wide association study of more than 40,000 Bipolar disorder cases provides new insights into the underlying Biology. Nat. Genet. (2021) doi:10.1038/s41588-021-00857-4.

5. Viguera, A. C., Tondo, L. & Baldessarini, R. J. Sex differences in response to lithium treatment. Am. J. Psychiatry 157, 1509–1511 (2000).

6. Tohen, M. et al. Olanzapine versus lithium in the maintenance treatment of Bipolar disorder: a 12-month, randomized, douBle-Blind, controlled clinical trial. Am. J. Psychiatry 162, 1281–1290 (2005).

7. BALANCE investigators and collaBorators et al. Lithium plus valproate comBination therapy versus monotherapy for relapse prevention in Bipolar I disorder (BALANCE): a randomised open-laBel trial. Lancet 375, 385–395 (2010).

8. Grof, P. et al. Is response to prophylactic lithium a familial trait? J. Clin. Psychiatry 63, 942–947 (2002).

9. Hou, L. et al. Genetic variants associated with response to lithium treatment in Bipolar disorder: a genome-wide association study. Lancet 387, 1085–1093 (2016).

10. Song, J. et al. Genome-wide association study identifies SESTD1 as a novel risk gene for lithium-responsive Bipolar disorder. Mol. Psychiatry 21, 1290–1297 (2016).

11. Mertens, J. et al. Differential responses to lithium in hyperexcitaBle neurons from patients with Bipolar disorder. Nature 527, 95–99 (2015).

12. Stern, S. et al. Neurons derived from patients with Bipolar disorder divide into intrinsically different suB-populations of neurons, predicting the patients’ responsiveness to lithium. Mol. Psychiatry 23, 1453–1465 (2018).

13. Stern, S. et al. A Physiological InstaBility Displayed in Hippocampal Neurons Derived From Lithium-Nonresponsive Bipolar Disorder Patients. Biol. Psychiatry 88, 150–158 (2020).

14. Santos, R. et al. Deficient LEF1 expression is associated with lithium resistance and hyperexcitaBility in neurons derived from Bipolar disorder patients. Mol. Psychiatry (2021) doi:10.1038/s41380-020-00981-3.

15. Senner, F., Kohshour, M. O., ABdalla, S., Papiol, S. & Schulze, T. G. The Genetics of Response to and Side Effects of Lithium Treatment in Bipolar Disorder: Future Research Perspectives. Front. Pharmacol. 12, 638882 (2021).

16. Wolter, J. M., Jimenez, J. A., Stein, J. L. & Zylka, M. J. ToxCast chemical liBrary screen identifies diethanolamine as an activator of Wnt signaling. BioRxiv 2021.02.15.430319 (2021) doi:10.1101/2021.02.15.430319.

17. Newport, D. J. et al. Lithium placental passage and oBstetrical outcome: implications for clinical management during late pregnancy. Am. J. Psychiatry 162, 2162–2170 (2005).

18. Poels, E. M. P., Bijma, H. H., GalBally, M. & Bergink, V. Lithium during pregnancy and after delivery: a review. Int J Bipolar Disord 6, 26 (2018).

19. Munk-Olsen, T. et al. Maternal and infant outcomes associated with lithium use in pregnancy: an international collaBorative meta-analysis of six cohort studies. Lancet Psychiatry 5, 644–652 (2018).

20. ABu-Taweel, G. M. Effects of perinatal exposure of lithium on neuro-Behaviour of developing mice offspring. Indian J. Exp. Biol. 50, 696–701 (2012).

21. Messiha, F. S. Lithium and the neonate: developmental and metaBolic aspects. Alcohol 3, 107–112 (1986).

22. Poels, E. M. P. et al. Long-term neurodevelopmental consequences of intrauterine exposure to lithium and antipsychotics: a systematic review and meta-analysis. Eur. Child Adolesc. Psychiatry 27, 1209–1230 (2018).

23. Giles, J. J. & Bannigan, J. G. Teratogenic and developmental effects of lithium. Curr. Pharm. Des. 12, 1531–1541 (2006).

24. Chen, G., Rajkowska, G., Du, F., Seraji-Bozorgzad, N. & Manji, H. K. Enhancement of hippocampal neurogenesis By lithium. J. Neurochem. 75, 1729–1734 (2000).

25. Zanni, G. et al. Lithium Accumulates in Neurogenic Brain Regions as Revealed By High Resolution Ion Imaging. Sci. Rep. 7, 40726 (2017).

26. Santarelli, L. et al. Requirement of hippocampal neurogenesis for the Behavioral effects of antidepressants. Science 301, 805–809 (2003).

27. Sahay, A. & Hen, R. Adult hippocampal neurogenesis in depression. Nat. Neurosci. 10, 1110–1115 (2007).

28. Perera, T. D. et al. Antidepressant-induced neurogenesis in the hippocampus of adult nonhuman primates. J. Neurosci. 27, 4894–4901 (2007).

29. Madsen, T. M. et al. Increased neurogenesis in a model of electroconvulsive therapy. Biol. Psychiatry 47, 1043–1049 (2000).

30. Aygün, N. et al. Brain-trait-associated variants impact cell-type-specific gene regulation during neurogenesis. Am. J. Hum. Genet. 108, 1647–1668 (2021).

31. Liang, D. et al. Cell-type-specific effects of genetic variation on chromatin accessiBility during human neuronal differentiation. Nat. Neurosci. (2021) doi:10.1038/s41593-021-00858-w.

32. Stein, J. L. et al. A quantitative framework to evaluate modeling of cortical development By neural stem cells. Neuron 83, 69–86 (2014).

33. Valvezan, A. J. & Klein, P. S. GSK-3 and Wnt Signaling in Neurogenesis and Bipolar Disorder. Front. Mol. Neurosci. 5, 1 (2012).

34. Hur, E.-M. & Zhou, F.-Q. GSK3 signalling in neural development. Nat. Rev. Neurosci. 11, 539–551 (2010).

35. Qu, Z., Sun, D. & Young, W. Lithium promotes neural precursor cell proliferation: evidence for the involvement of the non-canonical GSK-3β-NF-AT signaling. Cell Biosci. 1, 18 (2011).

36. Nolen, W. A. et al. ISBD/IGSLI Task Force on the treatment with lithium What is the optimal serum level for lithium in the maintenance treatment of Bipolar disorder? A systematic review and recommendations from the ISBD/IGSLI Task Force on treatment with lithium Version 2. Bipolar Disord. 21, 394–409 (2019).

37. Kang, H. M. et al. Variance component model to account for sample structure in genome-wide association studies. Nat. Genet. 42, 348–354 (2010).

38. Nyholt, D. R. A simple correction for multiple testing for single-nucleotide polymorphisms in linkage disequiliBrium with each other. Am. J. Hum. Genet. 74, 765–769 (2004).

39. JerBer, J. et al. Population-scale single-cell RNA-seq profiling across dopaminergic neuron differentiation. Nat. Genet. 53, 304–312 (2021).

40. Wu, Y. et al. Colocalization of GWAS and eQTL signals at loci with multiple signals identifies additional candidate genes for Body fat distriBution. Hum. Mol. Genet. 28, 4161–4172 (2019).

41. The Schizophrenia Working Group of the Psychiatric Genomics Consortium, Ripke, S., Walters, J. T. R. & O’Donovan, M. C. Mapping genomic loci prioritises genes and implicates synaptic Biology in schizophrenia. BioRxiv (2020) doi:10.1101/2020.09.12.20192922.

42. Savage, J. E. et al. Genome-wide association meta-analysis in 269,867 individuals identifies new genetic and functional links to intelligence. Nat. Genet. 50, 912–919 (2018).

43. Meng, L. et al. Nucleostemin deletion reveals an essential mechanism that maintains the genomic staBility of stem and progenitor cells. Proc. Natl. Acad. Sci. U. S. A. 110, 11415–11420 (2013).

44. Tsai, R. Y. L. & McKay, R. D. G. A nucleolar mechanism controlling cell proliferation in stem cells and cancer cells. Genes Dev. 16, 2991–3003 (2002).

45. Meng, Q. et al. Integrative analyses prioritize GNL3 as a risk gene for Bipolar disorder. Mol. Psychiatry (2020) doi:10.1038/s41380-020-00866-5.

46. GTEx Consortium et al. Genetic effects on gene expression across human tissues. Nature 550, 204–213 (2017).

47. Thakore, P. I. et al. Highly specific epigenome editing By CRISPR-Cas9 repressors for silencing of distal regulatory elements. Nat. Methods 12, 1143–1149 (2015).

48. Pickar-Oliver, A. & GersBach, C. A. The next generation of CRISPR-Cas technologies and applications. Nat. Rev. Mol. Cell Biol. 20, 490–507 (2019).

49. Umans, B. D., Battle, A. & Gilad, Y. Where Are the Disease-Associated eQTLs? Trends Genet. (2020) doi:10.1016/j.tig.2020.08.009.

50. ForsBerg, L. et al. Maternal mood disorders and lithium exposure in utero were not associated with poor cognitive development during childhood. Acta Paediatr. 107, 1379–1388 (2018).

51. Major, M. B. et al. Wilms tumor suppressor WTX negatively regulates WNT/Beta-catenin signaling. Science 316, 1043–1046 (2007).

52. CampBell, K. Dorsal-ventral patterning in the mammalian telencephalon. Curr. Opin. NeuroBiol. 13, 50–56 (2003).

53. Hallonet, M. et al. Vax1 is a novel homeoBox-containing gene expressed in the developing anterior ventral foreBrain. Development 125, 2599–2610 (1998).

54. Jun, G. et al. Detecting and estimating contamination of human DNA samples in sequencing and array-Based genotype data. Am. J. Hum. Genet. 91, 839–848 (2012).

55. Malek, M. et al. flowDensity: reproducing manual gating of flow cytometry data By automated density-Based cell population identification. Bioinformatics 31, 606–607 (2015).

56. Kang, H. M. et al. Efficient control of population structure in model organism association mapping. Genetics 178, 1709–1723 (2008).

57. Fairley, S., Lowy-Gallego, E., Perry, E. & Flicek, P. The International Genome Sample Resource (IGSR) collection of open human genomic variation resources. Nucleic Acids Res. 48, D941–D947 (2020).

58. Civelek, M. et al. Genetic Regulation of Adipose Gene Expression and Cardio-MetaBolic Traits. Am. J. Hum. Genet. 100, 428–443 (2017).

59. Devlin, B. & Roeder, K. Genomic control for association studies. Biometrics 55, 997–1004 (1999).

60. Wolter, J. M. et al. Cas9 gene therapy for Angelman syndrome traps UBe3a-ATS long non-coding RNA. Nature (2020) doi:10.1038/s41586-020-2835-2.

61. 1000 Genomes Project Consortium et al. A gloBal reference for human genetic variation. Nature 526, 68–74 (2015).

62. MatoBa, N. et al. Common genetic risk variants identified in the SPARK cohort support DDHD2 as a candidate risk gene for autism. Transl. Psychiatry 10, 265 (2020).

